# Dynamic acylome reveals metabolite driven modifications in *Syntrophomonas wolfei*

**DOI:** 10.1101/2022.07.18.500474

**Authors:** Janine Y. Fu, John M. Muroski, Mark A. Arbing, Jessica A. Salguero, Neil Q. Wofford, Michael J. McInerney, Robert P. Gunsalus, Joseph A. Loo, Rachel R. Ogorzalek Loo

**Author notes:** **Correspondence:** Rachel R. Ogorzalek Loo.

## Abstract

*Syntrophomonas wolfei* is an anaerobic syntrophic microbe that degrades short-chain fatty acids to acetate, hydrogen, and/or formate. This thermodynamically unfavorable process proceeds through a series of reactive acyl-Coenzyme A species (RACS). In other prokaryotic and eukaryotic systems, the production of intrinsically reactive metabolites correlates with acyl-lysine modifications, which have been shown to play a significant role in metabolic processes. Analogous studies with syntrophic bacteria, however, are relatively unexplored and we hypothesize that highly abundant acylations could exist in *S. wolfei* proteins, corresponding to the RACS derived from degrading fatty acids. Here, by mass spectrometry-based proteomics (LC-MS/MS), we characterize and compare acylome profiles of two *S. wolfei* subspecies grown on different carbon substrates. Because modified *S. wolfei* proteins are sufficiently abundant for post-translational modification (PTM) analyses without antibody enrichment, we could identify types of acylations comprehensively, observing six types (acetyl-, butyryl-, *3-*hydroxybutyryl-, crotonyl-, valeryl-, hexanyl-lysine), two of which have not been reported in any system previously. All of the acyl-PTMs identified correspond directly to RACS in fatty acid degradation pathways. A total of 369 sites of modification were identified on 237 proteins. Changing the carbon substrate altered the acylation profile. Moreover, structural studies and *in vitro* acylation assays of a heavily modified enzyme, acetyl-CoA transferase, provided insight on the possible impact of these acyl-protein modifications. Our findings link protein acylation by RACS to shifts in cellular metabolism.

## 1 Introduction

Microbial syntrophy is an important component of the global carbon cycle. Recycling carbon anaerobically requires cooperation and cross-feeding between different microbial species to decompose large biopolymers initially into sugars, amino acids, aromatic acids, which are then fermented to longer chain fatty acids and alcohols (McInerney, Sieber & Gunsalus, 2009). Syntrophic bacteria play a key role in anaerobic decomposition by degrading the fatty acids and alcohols produced by fermentative bacteria to acetate, hydrogen, formate, and CO2, which are substrates for their methanogenic partners (Schink, 1997). Metabolic cooperation during syntrophic metabolism is essential due to the unfavorable thermodynamics of these reactions if syntrophic end products such as hydrogen and formate accumulate. Under standard conditions, the degradation of aliphatic and aromatic acid intermediates by syntrophs is endergonic and the associated free energy is highly dependent on entropy (McInerney & Beaty, 1988). In effect, the flux of compounds into and out of the system plays a large role in whether reactions will proceed forward spontaneously. By rapidly consuming the products of fatty acid degradation (hydrogen, formate, and acetate), methanogens keep the concentrations of these compounds low enough for syntrophic fatty acid degradation to become exergonic (McInerney & Beaty, 1988). The result is a tightly knit network of interacting microbial species that rely on one another in order to convert biopolymers to methane and carbon dioxide. A thorough understanding of the metabolic interactions during the conversion of organic carbon to methane, especially syntrophy, is key to understanding the fate of organic carbon in anaerobic environments (McInerney, Sieber & Gunsalus, 2009; Stams et al., 2012; Boll et al., 2020; Amos & McInerney, 1989).

*Syntophomonas wolfei* is a syntrophic bacterium that is a leading model for examining how microbial communities degrade short-chain fatty acids (McInerney et al., 1981; Lorowitz, Zhao & Bryant, 1989). To metabolize saturated fatty acids, the organism must be co-cultivated with a hydrogen/formate-consumer; typically the model methanogen *Methanospirillum hungatei* has been used (McInerney, Bryant & Pfennig, 1979; McInerney et al., 1981). However *S. wolfei* can degrade unsaturated fatty acids, such as crotonate, in either pure or coculture (Beaty & McInerney, 1987; Amos & McInerney, 1990), thus enabling comparisons of how this bacterium shifts its carbon metabolism to adapt to changing carbon substrates. The ability to grow *S. wolfei* in pure and multispecies cultures has resulted in its use as a model for studying anaerobic, syntrophic butyrate degradation. *S. wolfei* utilizes β-oxidation to degrade short chain fatty acids, for which the *S. wolfei* subspecies *wolfei* strain Göttingen genome encodes many paralogs: nine acyl-CoA dehydrogenase genes, five enoyl-CoA hydratase genes, six *3-*hydroxyacyl-CoA dehydrogenase genes, and five acetyl-CoA acetyltransferase genes (Sieber et al., 2010). While the main enzymatic steps have been identified, many questions remain about substrate specificity and regulation of these paralogs. Proteomic evidence indicates that, remarkably, altering growth from axenic (cultivation as a single species) to syntrophic conditions (cultivation with *M. hungatei)* does not change protein abundance significantly (Sieber et al., 2015; Crable et al., 2016; Schmidt et al., 2013). Despite the consistency in enzymatic constituents, it is evident that enzymatic catalysis rates do change with growth condition (McInerney & Wofford, 1992; Wofford, Beaty & McInerney, 1986). If these activity changes do not reflect shifts in cellular transcription or translation, we hypothesized that the enzymes themselves were impacted by post-translational modifications (PTMs) in a manner that correlates with the changing environment.

PTMs are present in all kingdoms of life and have a plethora of structures, physicochemical properties, and by extension, functions (Walsh & Jefferis, 2006). PTMs have long been attractive candidates for regulating various cellular functions and metabolic pathways; intracellular conditions can be sensed and directly alter proteins. One class of such PTMs is lysine acylation, a family of diverse modifications. Uniquely, lysine can be acylated either by enzyme-mediated reactions (Ali et al., 2018) or spontaneously by reactive metabolites like reactive acyl-Coenzyme A species (RACS) or acyl phosphates (Wagner & Hirschey, 2014; Wagner & Payne, 2013; Baeza, Smallegan & Denu, 2015). The resulting acylation events have been shown to both impede and initiate enzymatic activity (Gardner et al., 2006; Garrity et al., 2007). These PTMs have been identified across all domains of life (Verdin & Ott, 2015; VanDrisse & Escalante-Semerena, 2019; Christensen et al., 2019) and have conserved mechanisms of acyl-regulation (Sanders, Jackson & Marmorstein, 2010; Greiss & Gartner, 2009). In eukaryotes, these modifications are primarily found in the mitochondria and are implicated in a wide range of human metabolic-related diseases (Anderson & Hirschey, 2012; Baldensperger & Glomb, 2021; Ali et al., 2018). In prokaryotes, acyl-PTMs modify proteins involved in metabolism in addition to other biological processes such as transcription, chemotaxis, and protein stability (VanDrisse & Escalante-Semerena, 2019; Christensen et al., 2019; Bernal et al., 2014).

Protein lysine acylation by RACS can result directly from the flux through the metabolic pathways that generate the reactive metabolites. These PTMs may record variations in carbon flux due to stress or changes in substrate use, providing a mechanism to regulate carbon flow depending on carbon flux conditions (Schilling et al., 2015; Trub & Hirschey, 2018). Acyl-lysine modifications were characterized recently in *Syntrophus aciditrophicus* and shown to correspond directly to RACS generated from the syntroph’s degradation of benzoate (Muroski et al., 2022). Enzymes involved in degrading benzoate were not only modified with the respective acyl-CoA intermediate utilized or produced, but also with additional RACS that are produced by other enzymes in the pathway. The fact that active deacylases were identified suggest acylations impact syntrophic metabolism. *S. aciditrophicus* displayed acylation levels abundant enough to forego antibody enrichment strategies that are often used to study protein modifications in mass spectrometry-based proteomics analyses (Zhao & Jensen, 2009; Mertins et al., 2013) and this study raised the possibility that abundant acylations exist in other syntrophs like *S. wolfei*, that generate RACS during their metabolism. If acyl-PTMs are a feature of syntrophy, they might play an important role in syntrophic metabolism, which would be indicated by changes in PTMs with carbon substrate.

Here, we use mass spectrometry-based proteomics to take a systems level approach to identify acyl-modifications in the *S. wolfei* proteome from cells cultivated on different environmental substrates. Without utilizing PTM-specific enrichment strategies, we demonstrate how the acylome profile changes with growth condition —qualitatively in the acyl groups that modify lysines and quantitatively in their abundance. In particular, enzymes involved in the β-oxidation pathway were found to be heavily decorated with a wide range of acyl groups, all of which are related to RACS found in fatty acid oxidation. Specific sites changed in both the type and abundance of modifications observed under different conditions. To probe the effect that PTMs may have on protein function, structural studies and *in vitro* acylation assays were performed on acetyl-CoA transferase, a β-oxidation pathway enzyme observed to be extensively modified.

## 2 Materials and methods

### 2.1 Culturing of cells

*Syntrophomonas wolfei* subspecies *wolfei* strain Göttingen (DSM 2245B) (Lorowitz, Zhao & Bryant, 1989) (hereafter referred to as *S. wolfei* Göttingen) cells were grown axenically with crotonate as the carbon substrate (McInerney et al., 1981), or in coculture with methanogenic partner, *Methanospirillum hungatei* JF1 (DSM864), on butyrate as previously described (McInerney, Bryant & Pfennig, 1979). Cells were harvested anaerobically and then frozen and stored at -70°C until processed (Muroski et al., 2022). *Syntrophomonas wolfei* subspecies *methylbutyratica* strain 4J5T (JCM 14075) was grown in the presence of methanogenic partner *M. hungatei* as described above but using the substrates, butyrate, crotonate, 2-methylbutyrate, valerate or hexanoate. Analyses were performed on three biological replicates from each culture condition.

### 2.2 Sample preparation

Cell pellets were suspended in 4.0% v/v ammonium lauryl sulfate, 0.1% w/v sodium deoxycholate, and 5 mM tris(2-carboxyethyl)phosphine in 100 mM ammonium bicarbonate (ABC) for lysis. Proteins were alkylated and digested using enhanced filter-assisted sample preparation (eFASP) as described by Erde et al (Erde, Loo & Loo, 2014). Briefly, the lysate was exchanged into buffer composed of 8 M urea, 0.1% w/v sodium deoxycholate, and 0.1% w/v *n*-octyl glucoside with a 10 kDa Microcon ultrafiltration unit (Millipore). Proteins were alkylated with 17mM iodoacetamide and digested overnight at 37°C with a 1:100 ratio of trypsin:protein. Detergents were extracted away from the peptide-containing buffer using ethyl acetate. Peptides were dried and stored at -20°C until used in MS experiments or for offline fractionation.

### 2.3 Mass spectrometry (MS)

Peptides were dried, resuspended in 0.1% acetic acid, desalted with STAGE tips fabricated from 3M Empore C18 solid phase extraction disks, and then re-dried (Rappsilber, Mann & Ishihama, 2007). Resultant peptide-containing samples were resuspended in LC-MS injection buffer (3% acetonitrile and 0.1% formic acid) and analyzed on an Orbitrap Exploris™ 480 Mass Spectrometer (ThermoFisher Scientific) using liquid chromatography-tandem mass spectrometry (LC-MS/MS). Chromatography employed a reverse phase EASY-Spray™ column (25cm x 75μm ID, PepMap™ RSLC C18, 2μm, ThermoFisher Scientific) connected to an UltiMate™ 3000 RSLCnano HPLC (ThermoFisher Scientific). Mobile phase buffers A (0.1% formic acid) and B (0.1% formic acid in 100% acetonitrile) were delivered at 300 nL/min with the following gradient: 3-20% B in 115 minutes, 20-32% B in 19 minutes, 32-95% B in 1 minute.

Data-dependent acquisition (DDA) mode was used to select ions for tandem MS. Positive ion precursor scans (375-1800 *m/z*) were acquired at 60,000 resolution with a normalized automatic gain control (AGC) target of 100%. Peptide ions were fragmented using higher-energy collisional dissociation (HCD) at a normalized collisional energy of 27%. Dynamic exclusion was applied for 45s over ±10 ppm. MS/MS scans were collected with a first fixed mass of *m/z* 100, 2 *m/z* isolation window, 15000 orbitrap resolution, and normalized AGC target of 100%.

### 2.4 Hydrophilic interaction liquid chromatography

To expand the number of tryptic peptides detected and establish a list of target peptide ions for parallel reaction monitoring (PRM) (Peterson et al., 2012) MS experiments, peptide products from eFASP were fractionated off-line by hydrophilic interaction chromatography (HILIC) (Alpert, 1990). Tryptic peptides (50 μg) were loaded onto a BioPureSPN MACRO PolyHYDROXYETHYL A™ column (The Nest Group) in 90% acetonitrile and 150 mM ammonium formate, pH 3. Peptides were then eluted in six fractions sequentially using: 80%, 78%, 74%, 71%, 40%, and 35.5% acetonitrile, respectively. Peptide eluates were dried, resuspended, and desalted with C18 STAGE tips as described above. For *S. wolfei* Göttingen only, fractions 1 and 6 were combined for analysis to reduce instrument operation time. *S. wolfei* subsp. *methylbutyratica* fractions 1 and 6 were analyzed independently to optimize detection of hydrophobic, long-chain acyl-PTMS in fraction 1. HILIC fractionated peptides were analyzed by LS-MS/MS as described above.

### 2.5 Acyl quantification

Unscheduled PRM-targeted MS analysis (Peterson et al., 2012) was utilized to quantify acyl-peptides of interest using the list of target peptide ions identified in HILIC DDA experiments. Desalted peptides from the eFASP procedure were analyzed without any offline fractionation. Liquid chromatography was performed identically to shotgun and HILIC-fractionated experiments. The inclusion list used for PRMs is shown in **Supplemental Table 1**. The MS2 scans were obtained at a 17500 resolution with a normalized AGC target 100% and an isolation window of *m/z* 1.2 using HCD fragmentation at a normalized collision energy of 30%.

**Table 1.**
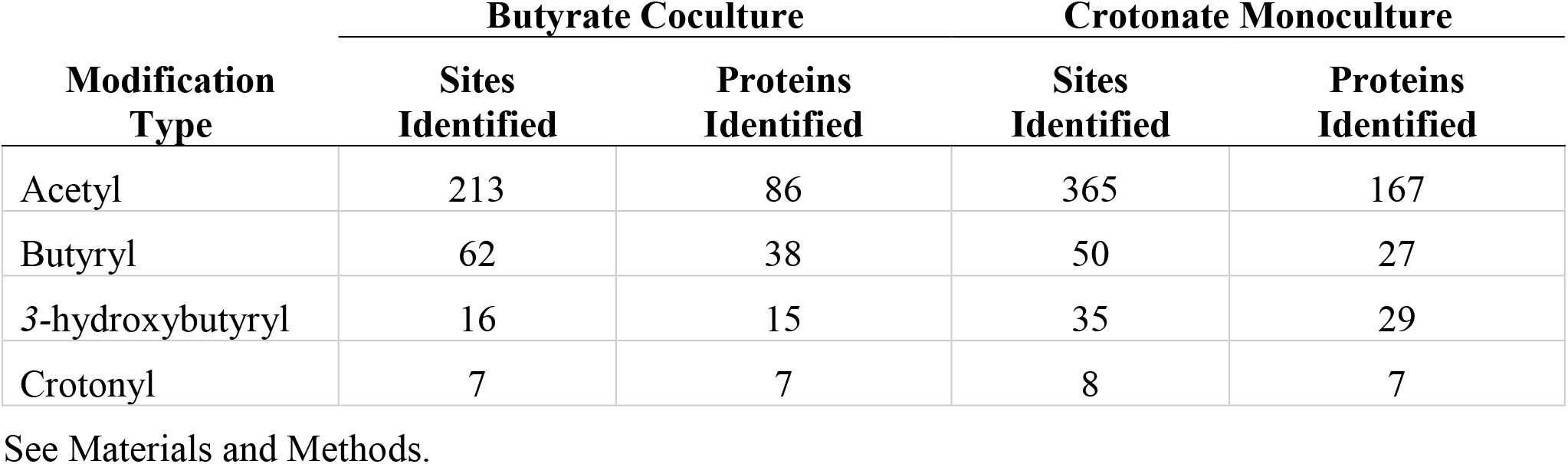
Overall acylation sites detected in *S. wolfei* Göttingen.

| Modification<br>Type | Butyrate Coculture |  | Crotonate Monoculture |  |
| --- | --- | --- | --- | --- |
|  | Sites<br>Identified | Proteins<br>Identified | Sites<br>Identified | Proteins<br>Identified |
| Acetyl | 213 | 86 | 365 | 167 |
| Butyryl | 62 | 38 | 50 | 27 |
| 3-hydroxybutyryl | 16 | 15 | 35 | 29 |
| Crotonyl | 7 | 7 | 8 | 7 |

See Materials and Methods.

### 2.6 Analysis of acyl-peptide standards to validate identifications

Synthetic acylated peptide standards, TPIGK(valeryl)FLGQFK and TPIGK(hexanyl)FLGQFK, (1 pmol each) were desalted with STAGE tips and delivered for tandem MS in DDA mode, using the same LC-MS/MS parameters as described above.

### 2.7 Data analysis

RAW data files from DDA experiments were analyzed using ProteomeDiscoverer™ (version 1.4) and database searched with the Mascot search algorithm (Matrix Science, version 2.6.2) (Perkins et al., 1999). UniProt *S. wolfei* Göttingen and *M. hungatei* protein sequences (as of February 1, 2017) were concatenated to sequences of common contaminants and searched against samples from the respective cell cultures. The total number of entries in this database is 5691. *S. wolfei* subsp. *methylbutyratica* protein sequences from the draft genome available at JGI (Narihiro et al., 2016) and UniProt *M. hungatei* protein sequences (as of April 7, 2020) were concatenated to sequences of common contaminants and searched against samples from the respective cell cultures. The total number of database entries is 6124. Parameters selected for the Mascot search were: enzyme name, trypsin; maximum missed cleavage sites, 2; precursor mass tolerance, 10 ppm; fragment mass tolerance, 0.02 Da; and variable methionine oxidation and cysteine carbamidomethylation. Searches also considered the potential acyl-lysine modifications listed in **Supplemental Table 2**. The selection of acyl modifications to be searched was based on RACS identified in the β-oxidation pathways of *S. wolfei* Göttingen and *S. wolfei* subsp. *methylbutyratica*. The requirement for protein identification is that at least 2 unique peptides are detected with a Mascot ion score ≥ 20, which corresponds to a 95% confidence, amongst the three biological replicates. Protein abundances were normalized by species to account for differences in cell pellet compositions between mono and cocultures. Proteomic datasets submitted to the ProteomeXchange Consortium through the MassIVE repository are identified as PXD034881.

For the targeted analyses (PRM experiments), ProteomeDiscoverer ™ search results from HILIC fractions were imported into Skyline (version 21.1.0.278, MacCoss lab, University of Washington) to build spectral libraries (Maclean et al., 2010). PRM RAW files were imported into Skyline for quantitative data processing and proteomic analysis. The protein sequence database of *S. wolfei* Göttingen described above was added to Skyline as the background proteome. Peak areas from extracted ion chromatograms for four to six coeluting product ions were selected for quantification based on a higher dot product correlation between observed transitions of target peptides and library spectrum, indicating higher confidence in peptide detection. Cyclized immonium ions (Muroski et al., 2021) for acyl-lysine modifications were added to the Skyline analysis as customized product ions. All integrated peaks were manually inspected to ensure correct peak detection and integration. Peak areas of selected product ions were summed for acyl-peptide quantification. As an internal control, the abundance of each acyl-peptide was normalized to the abundance of an unmodified peptide in the same protein.

### 2.8 Protein expression and purification of *S. wolfei* Göttingen Act2 (Swol_0675)

The Swol_0675 gene was synthesized and cloned into pMAPLe4 (Arbing et al., 2013) (Twist Bioscience) which appends an N-terminal tag encoding a hexahistidine tag and TEV protease cleavage site to the target protein. The expression plasmid was transformed into *E. coli* BL21-Gold (DE3) and an overnight inoculum was used to inoculate 2L of Terrific broth supplemented with kanamycin. Cultures were grown at 37°C and protein expression was induced by the addition of IPTG to 0.5 mM when the culture reached an OD600 of 1.0; growth was continued at 18°C overnight and the cells harvested by centrifugation approximately 18h after induction. The cell pellet was resuspended in Buffer A (50 mM Tris pH 8.0, 300 mM NaCl, 20 mM imidazole, 5 mM β-mercaptoethanol) supplemented with EDTA (1 mM), PMSF (1 mM), and benzonase. The cell suspension was lysed using an Emulsiflex C-3 and the lysate clarified by centrifugation (39kxG, 40’). The clarified lysate was loaded on a 5 mL NiNTA column equilibrated in Buffer A and the column was washed extensively with buffer A before eluting bound protein with a linear gradient of Buffer B (Buffer A with 300 mM imidazole). TEV protease was added to the peak fractions and the digest was dialyzed against Buffer A overnight. The following day the digest was passed over the NiNTA column to remove H6-tagged TEV protease and the cleaved affinity tag and the flow-through were collected. The flow-through was concentrated and further purified by size exclusion chromatography using a Superdex 200 equilibrated with 20 mM Tris pH 8.0, 150 mM NaCl, 5 mM β-mercaptoethanol. The peak containing Act2 was concentrated using an Amicon Ultra-15 centrifugal filter unit (10 kDa MWCO) to 30 mg/mL.

### 2.9 Crystallization and structure determination of *S. wolfei* Göttingen Act2 (Swol_0675)

Crystals of Act2 were grown at room temperature using the hanging drop vapor diffusion method with 0.1 M sodium formate pH 7.0, 12% PEG 3350 as the reservoir solution. Crystals were cryoprotected with a brief soak in 23% PEG 3350, 5% PEG 400 before being flash frozen. Diffraction data was collected at the Northeastern Collaborative Access Team (NE-CAT) facility, beamline 24-ID-C, at the Advanced Photon Source at Argonne National Laboratory. Data were processed with XDS(Kabsch & IUCr, 2010) and the structure solved by molecular replacement using the program phenix.phaser (McCoy et al., 2007) with a search model generated by the Phyre2 server.(Kelley et al., 2015) The model was refined with phenix.refine. (Afonine et al., 2012) Crystallographic data and refinement statistics are listed in **Table 4** (prepared with phenix.table_one) and the structure has been deposited in the Protein Data Bank under accession code 7N7Z.

### 2.10 *In vitro* acylation assay

Purified *S. wolfei* Göttingen Swol_0675 recombinant protein (Act2, 5μM) was incubated either with 1 μM acetyl-CoA or 50 μM butyryl-CoA in 100mM ABC for 1 or 3 hour(s), respectively, at 37°C as described previously (Parks & Escalante-Semerena, 2020). Reactions were stopped by exchanging into 100mM ABC buffer to remove excess acetyl-CoA using 10 kDa MWCO Amicon® centrifugal filters (Millipore). Modified proteins were then digested for analysis by LC-MS/MS. Briefly, the modified proteins were heated to 95°C for 10 min, disulfides were reduced with 20mM dithiothreitol (DTT) for 1hr at 60°C, and alkylated by incubation with 50mM iodoacetamide at room temperature in the dark for 45 min. Following incubation, iodoacetamide was quenched by adding excess DTT. Acylated Swol_0675 was digested overnight with trypsin (1:100, 37°C), followed by a second overnight digest with endoproteinase GluC (1:100, 25°C). Peptides were desalted with STAGE tips. The mass spectrometer was operated in DDA mode, as described earlier, with the LC gradient: 3-20% B in 32 minutes, 20-32% B in 5 minutes, 32-95% B in 1 minute. MS data was analyzed as described above.

## 3 Results

### 3.1 Comparing the proteomes of butyrate and crotonate grown *S. wolfei* Göttingen

*S. wolfei* Göttingen metabolizes butyrate as shown in **Figure 1**, relying on methanogen *M. hungatei* to maintain H2 and/or formate levels low enough so that butyrate degradation is thermodynamically favorable. *S. wolfei* does not require a partner organism to metabolize crotonate, because for every molecule of crotonate oxidized down the pathway, another is reduced to butyrate to consume the electrons generated by crotonate oxidation. Whole cell extracts from *S. wolfei* Göttingen grown on crotonate in pure culture or on butyrate in coculture with *M. hungatei* were trypsin-digested and analyzed by tandem mass spectrometry to examine how carbon substrates influenced protein abundances. Overall abundance differences between the two carbon substrates are illustrated in **Figure 1A**. Out of the 446 proteins identified in *both* the crotonate monoculture (CM) and butyrate coculture (BC), only 71 proteins had significant changes in protein abundance (>2x fold change, p<0.05), with 57 and 14 proteins up-regulated in CM and BC cultures, respectively. Out of these 71 proteins, 39 proteins had fold changes greater than 3-fold, and many of these changes were relatively modest mirrors of observations in previous proteomic studies (Sieber et al., 2015; Crable et al., 2016; Schmidt et al., 2013).

**Figure 1.**
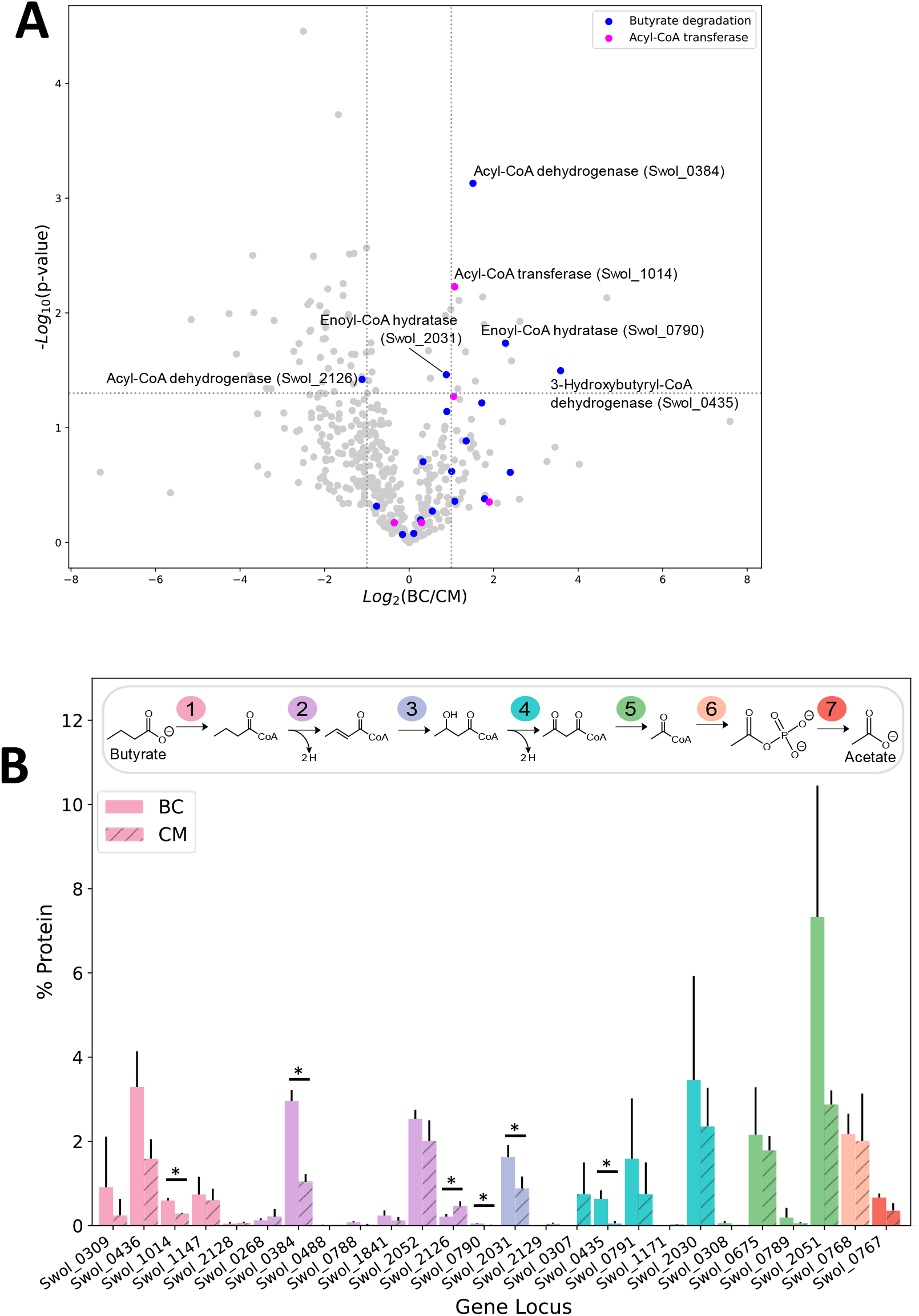
Protein abundance of *S. wolfei* Göttingen proteome between carbon substrates. **(A)** Volcano plot of protein-fold changes between butyrate cocultures (BC) and crotonate monocultures (CM). Dotted lines represent significance cutoffs (p < 0.05, fold-change >2 or < 0.5) **(B)** Percent protein abundance of beta-oxidation paralogs. Colors indicate enzymes with multiple paralogs. Each number in pathway represents the step catalyzed by the paralogs. (*p<0.05)

Our MS analysis was able to achieve greater coverage of the proteome, identifying proteins not previously detected. Genomic and proteomic analysis has showed that *S. wolfei* Göttingen contains and simultaneously expresses multiple paralogs for each step in β-oxidation (Sieber et al., 2010, 2015). Improved MS technologies enabled our proteomic analyses to detect 15 additional low-abundance paralogs that were not seen in previous work **(Figure 1B)** (Sieber et al., 2015; Crable et al., 2016; Schmidt et al., 2013). These identified proteins encompass the first five steps in β-oxidation and are as follows: acyl-CoA transferase (Swol_0309, 1014, 1147, 2128), acyl-CoA dehydrogenase (Swol_0488, 0788, 1841), enoyl-CoA hydratase (Swol_0790, 2031, 2129), *3-*hydroxybutyryl-CoA dehydrogenase (Swol_0307, 0791, 1171), and acetyl-CoA acetyltransferase (Swol_0308, 0789). Our improved proteomic depth enabled detection of additional β-oxidation paralogs and led us to investigate the presence of protein acyl-modifications which have been largely unexplored in *S. wolfei*.

### 3.2 The *S. wolfei* Göttingen proteome has a wide range of acyl-lysine modifications

A study in another syntroph, *S. aciditrophicus*, identified several types of acylations related to the RACS generated in the microbe’s metabolism (Muroski et al., 2022). This investigation raised the possibility that acylations could be similarly abundant in other syntrophs, like *S. wolfei* Göttingen, which also produces RACS. To comprehensively identify acyl-lysine modifications in *S. wolfei* arising from RACS, we performed shotgun proteomics *without utilizing PTM-specific enrichment*. Protein lysates were also pre-fractionated by hydrophilic interaction chromatography (HILIC) to achieve greater depth in analyzing the acylome. Database searches incorporated mass shifts corresponding to putative modifications arising from RACS formed during butyrate and crotonate degradation: acetyl-, butyryl-, crotonyl, *3-*hydroxybutyryl, and acetoacetyl-lysine **(Supplemental Table S2)**. Of the 5 types of acylation predicted, we identified 4 (acetyl-, butyryl-, crotonyl-, and *3-*hydroxybutyryl-lysine) across both culture conditions. A total of 238 sites were identified in 111 proteins in BC grown cells and a total of 226 sites in 176 proteins in CM grown cells. These are large numbers for un-enriched samples. For comparison, enrichment for acetylated peptides from MV4-11 cells, a human acute myeloid leukemia cell line, using an antibody against acetyl-lysine identified 3600 acetylation sites on 1750 proteins, and experiments without antibody enrichment resulted in a 60-fold decrease in acetylation sites (Choudhary et al., 2009).

Of the acyl-PTMs identified in *S. wolfei* Göttingen, acetylation was most common, followed by butyrylation, *3-*hydroyxbutyrylation, and crotonylation, respectively (**Table 1**). Some lysine residues were identified with a variety of acyl-modifications; *e*.*g*., K159 from acyl-CoA transferase (Swol_1014) was observed with either an acetyl- or butyryl-lysine modification (**Table 2**). These two PTMs are each counted towards the number of sites identified for the respective modification type, but the lysine residue is only counted once in the total number of acyl-lysine sites identified. The type of acylation observed varies with culture condition, as displayed in **Figure 2A**. The number of unique acyl-peptides, a metric commonly reported in MS proteomics, encompasses peptides that contain multiple modified lysines, providing additional context that would be otherwise missed from reporting individual modified sites. **Supplemental Table 3** lists the modified peptide sequences detected. An important caveat is that the modification differences with carbon substrate may simply reflect interference from the *M. hungatei* proteins present in butyrate cocultures, reducing proteomic coverage of *S. wolfei*. To gauge this effect, we compared the number of *S. wolfei* proteins identified from the HILIC-fractionated butyrate cultures to those from crotonate. While 12% more *S. wolfei* proteins were identified from crotonate monocultures (981 vs. 873), a majority (768) of proteins were identified from both culture conditions, suggesting that the acylation differences observed primarily reflect the carbon substrate **(Supplemental Figure 1)**. Notably, all types of acyl-PTMs are present in both conditions, as is evident with butyryl-lysine identified in crotonate cultures. Butyryl-CoA, the respective RACS for butyrylation, is known to form in crotonate grown cells from the reduction of crotonyl-CoA during crotonate fermentation (McInerney, Sieber & Gunsalus, 2009). Observation of butyryl modification is also consistent with the fact that syntrophs have certain enzymatic steps in β-oxidation that are reversible (James et al., 2019). Indeed, the capability of *S. wolfei* to dismutate crotonate to acetate and butyrate is key to its axenic cultivation (McInerney & Beaty, 1988).

**Table 2.**
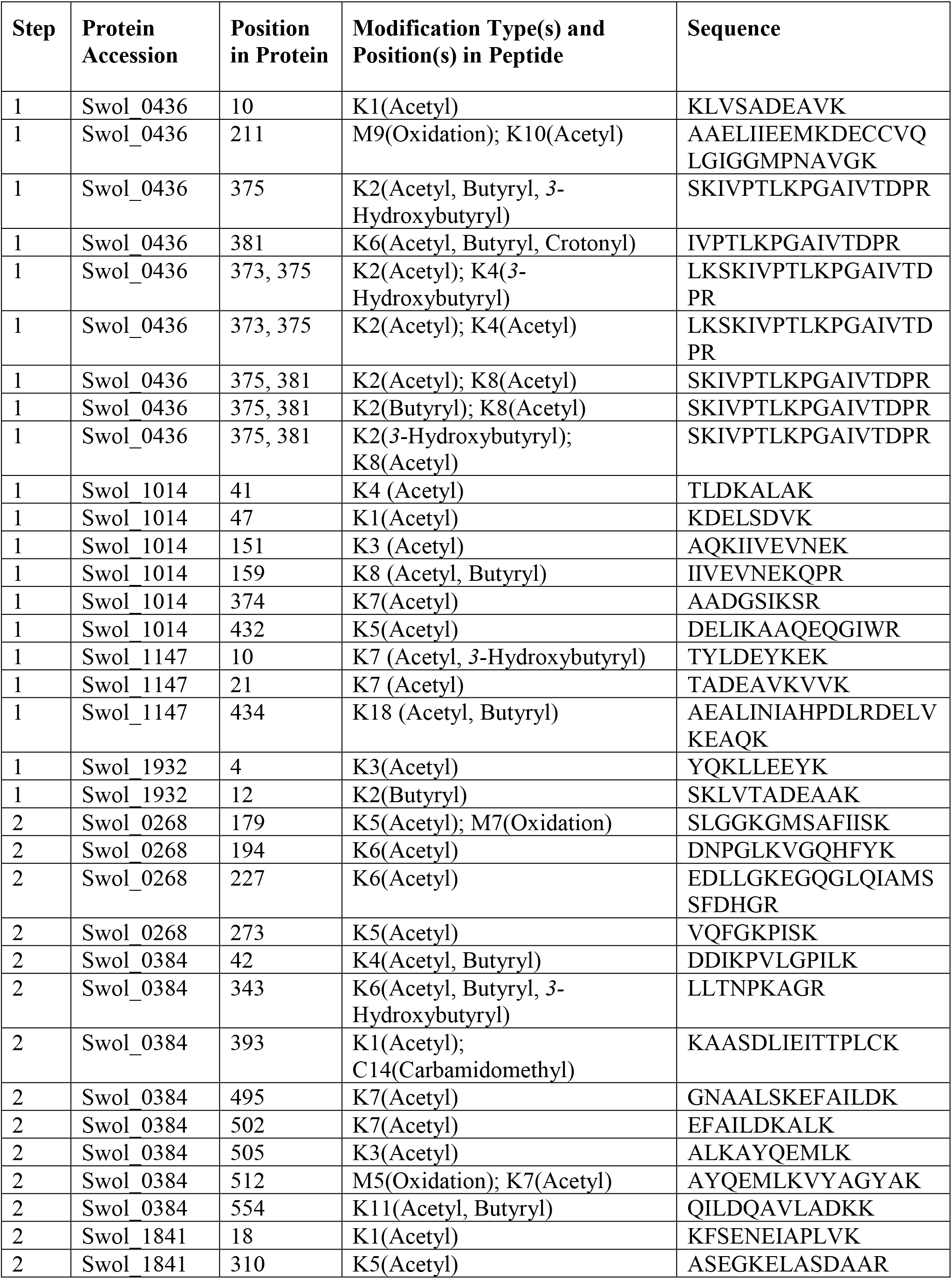
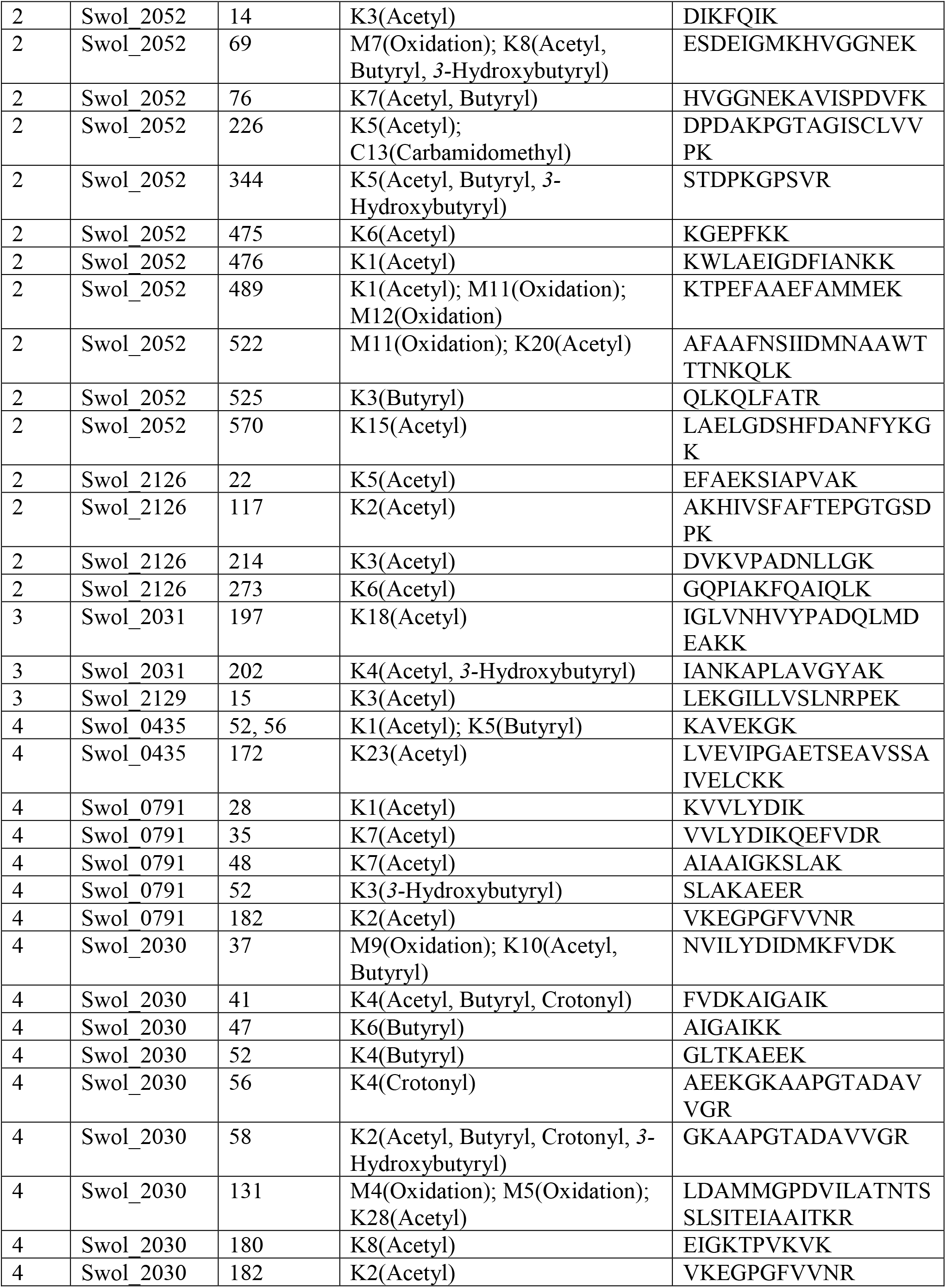
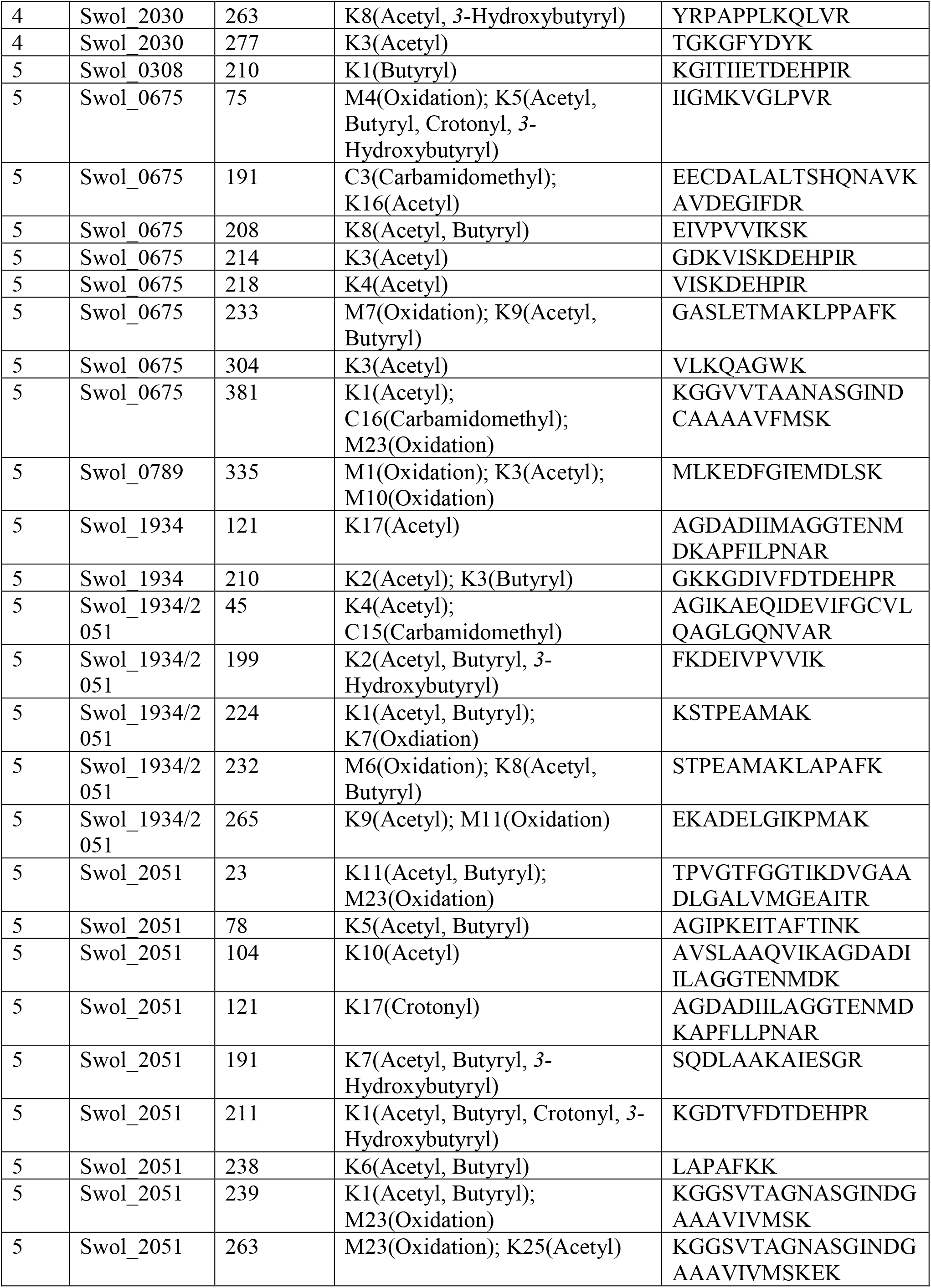
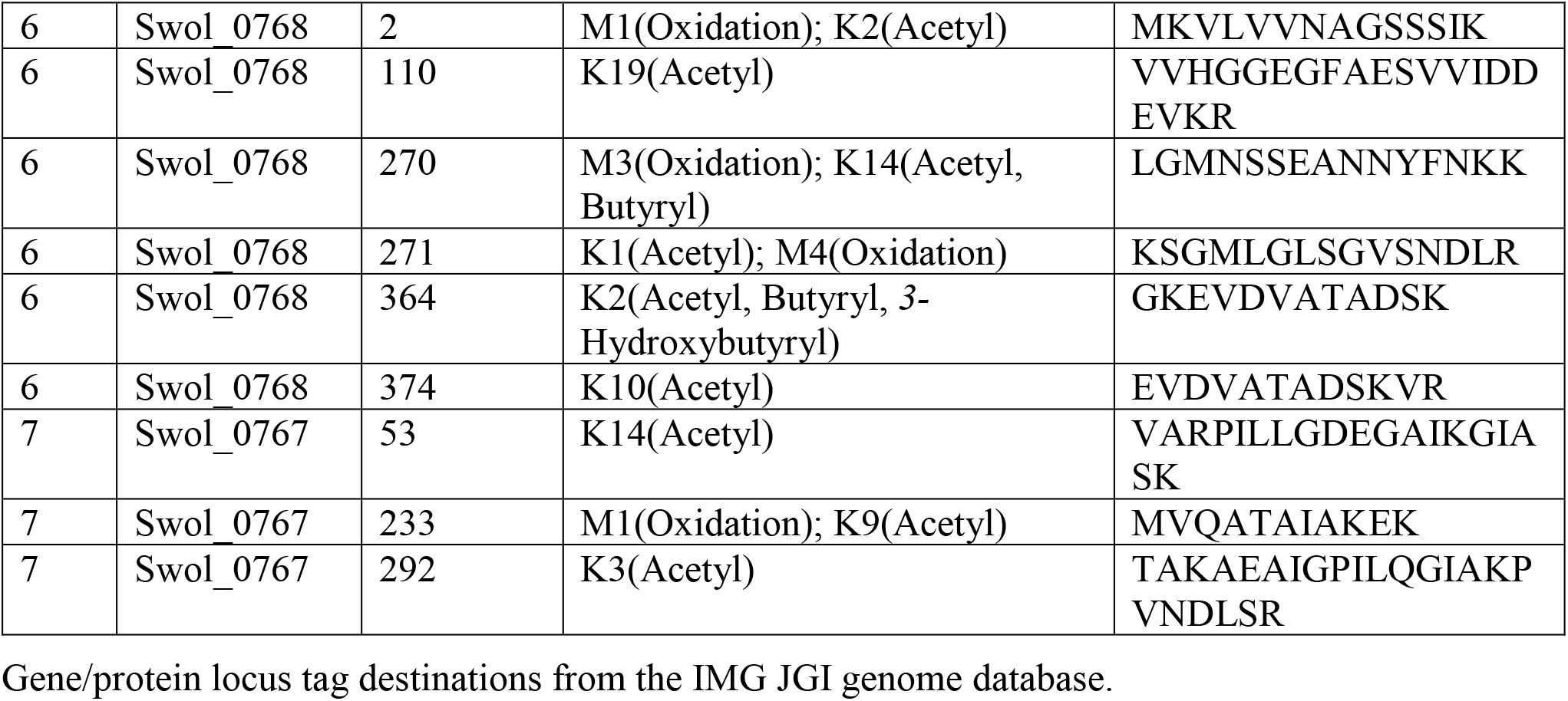
Acylation sites in β-oxidation pathway enzymes of *S. wolfei* Göttingen.

**Figure 2.**
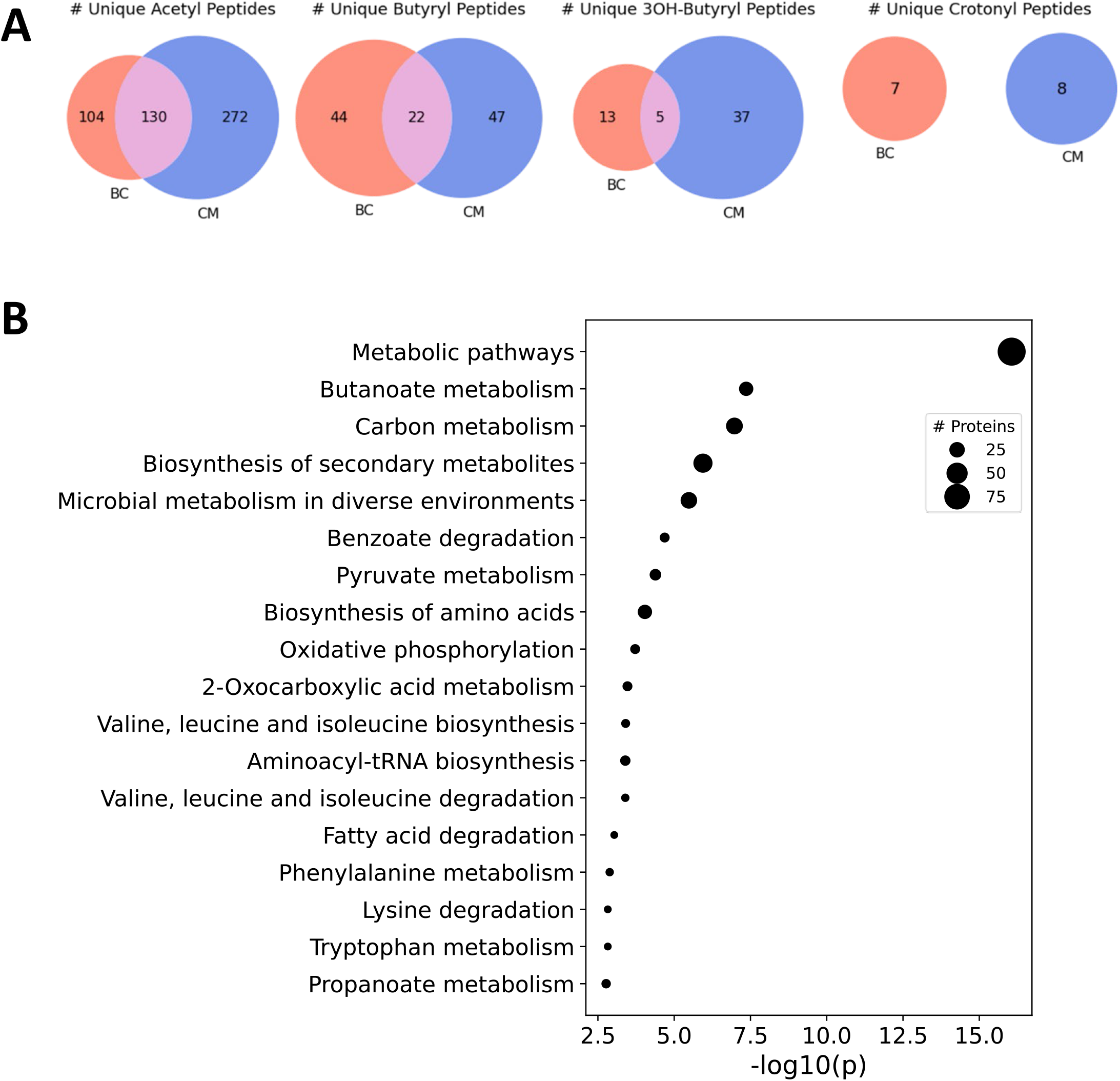
Acyl-lysine modifications identified across culture conditions in *S. wolfei* Göttingen. **(A)** Unique acyl-peptides identified between growth conditions BC and CM. **(B)** Significantly enriched pathways (KEGG Ontology) for the acylated proteins identified across both culture conditions. Point size illustrates the number of proteins identified.

### 3.3 Enzymes involved in *S. wolfei* Göttingen β-oxidation are extensively acylated

To understand the functional impact of these acyl-PTMs, a KEGG (Kyoto Encyclopedia of Genes and Genomes) pathway enrichment analysis was performed on the modified proteins (Kanehisa & Goto, 2000; Kanehisa & Sato, 2020). The most enriched processes were involved in metabolism, including the general categories butanoate metabolism, carbon metabolism, biosynthesis of secondary metabolites, and microbial metabolism in diverse environments **(Figure 2B)**. Indeed, enzymes involved in steps 1,2,4, and 5 of the butyrate β-oxidation pathway were heavily acylated, with 175 unique modified peptides identified on 21 out of the 26 proteins **(Table 2)**. In comparison, enzymes catalyzing glycolysis and gluconeogenesis had fewer sites of acylation; 9 unique modified peptides were identified on 5 out of the 26 proteins **(Supplemental Table 3)**. As there seemed to be a relationship between metabolic pathways producing RACS and protein acylation, we focused on the butyrate β-oxidation pathway. β-oxidation paralogs were grouped by the reaction catalyzed to discern trends in the number and type of acyl modifications identified. **Figure 3** shows that populations of enzymes involved in all of the catalytic steps are modified, with the largest number of unique, modified peptides found for acetyl-CoA transferase and the smallest number found for enoyl-CoA hydratases and acetate kinase. For the enzymes involved in β-oxidation, acetylation is the primary acylation for each of the steps in the pathway, followed by butyryl-, *3-*hydroxybutyryl-, and crotonyl-lysine modification. The low number of crotonyl-lysine modified peptides detected could be due to the very high activity of enoyl-CoA hydrolase in *S. wolfei* (McInerney & Wofford, 1992). This enzyme, which catalyzes the third step in β-oxidation, converts crotonyl-CoA to *3-*hydroxybutyryl-CoA and its high activity generates a smaller pool of crotonyl-CoA as compared to that of other RACS. The relative abundance of each type of acylation could be related to the intracellular concentrations of the respective acyl-CoA. The relative abundance of each acyl modification for enzymes involved in β-oxidation is similar to the acylation trends observed among all acylated proteins (those involved in other biological processes besides β-oxidation) in the *S. wolfei* proteome.

**Figure 3.**
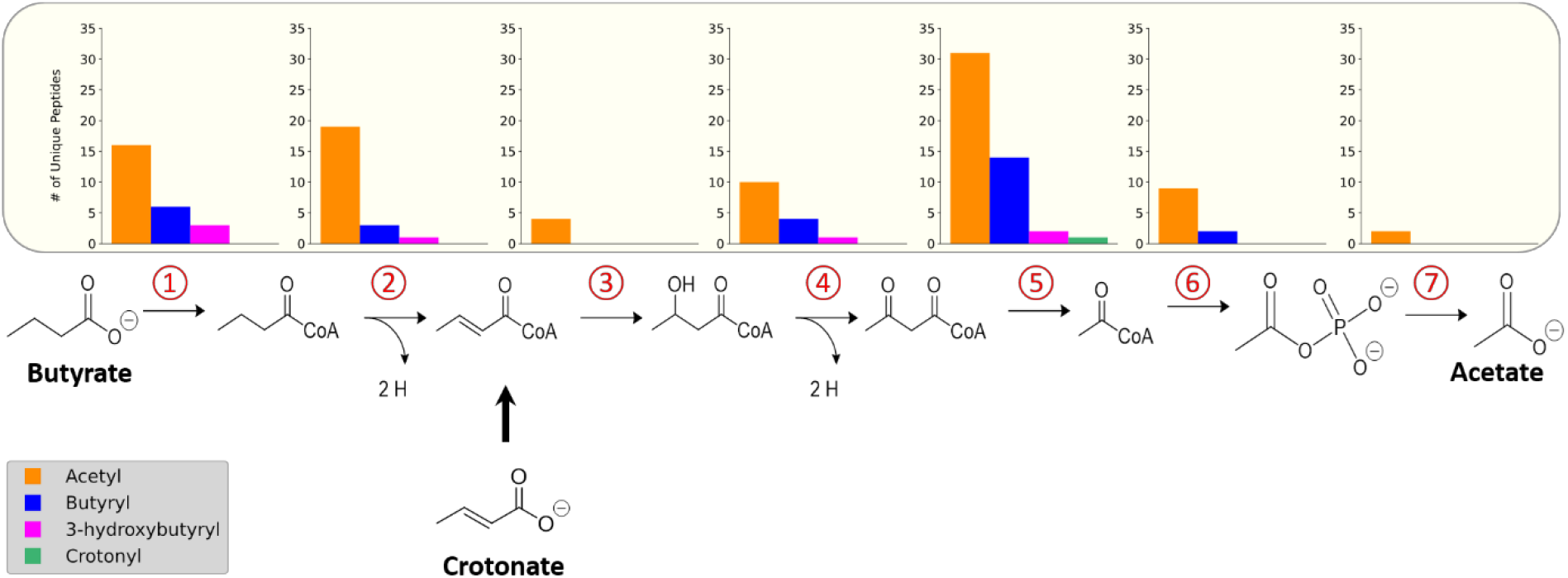
Acylated proteins identified in butyrate degradation pathway of *S. wolfei* Göttingen. Number of unique acylated peptides identified for each step in the pathway. Each number represents the step in the pathway catalyzed by the paralogs.

Many of the proteins involved in butyrate degradation that were similarly abundant in BC and CM grown cultures were found modified by several acyl-PTMs. Although proteins with significant abundance changes were also observed to be modified, these constitutively expressed proteins, Swol_0436, Swol_2052, Swol_2030, and Swol_2051 from steps 1,2,4, and 5, respectively, are of special interest. These proteins **(Table 2)** were particularly abundant compared to paralogs of the respective enzyme, maintained relatively stable expression (abundance) levels across growth conditions, and exhibited acylated residues. The presence of modifications on these proteins could suggest that post-translational regulation was in play.

### 3.4 Quantitation of *S. wolfei* Göttingen acyl-peptide abundances between cell growth conditions

In addition to investigating the presence, type, and number of modified sites, we next sought to measure the relative abundance of these acyl-lysine modifications in butyrate- and crotonate-grown cells. Targeted MS experiments, known as parallel reaction monitoring (PRM), were performed on the acyl-lysine peptides listed in **Table 2**. In PRM, the mass spectrometer detects ions for a user-selected peptide at a defined mass-to-charge ratio, often within a specified time interval, and the instrument isolates this precursor ion for fragmentation. This process enables acquisition of tandem mass spectra across the peptide’s elution period, yielding fragment ion chromatograms from which abundance is determined, allowing for a highly specific and selective MS approach for targeted peptide quantitation (Peterson et al., 2012). One of the key challenges in quantitatively comparing proteins from microbial consortia is that the relative distribution of organisms comprising the population (*e*.*g*., co-versus mono-culture) influences peptide abundance measurements (Chignell et al., 2018). To overcome this limitation, unmodified peptides from the respective acylated proteins were used as internal standards to normalize acylated peptide levels. Due to some modified peptides appearing in only one of the two culture conditions and peptide ionization efficiency limitations, not all the acyl-peptides could be quantified. The library of selected ions employed to quantify the selected peptides is shown in **Supplemental Table 1**. Targeted PRM capability is critical to quantify and localize PTMs, especially in peptides containing multiple modifications and for peptides that can be modified by different acyl groups. MS/MS level quantitation from PRM provides information on individual product ions, allowing confident discrimination between contributions from acyl-peptide variants. The utility of incorporating immonium ion derivatives for improved quantitation of acyl-peptides in PRM analyses is illustrated in **Supplemental Figure 2** (Muroski et al., 2021).

Acyl-peptide abundances differ in butyrate and crotonate grown cells. The data is complex, but trends can be identified. A majority of the acetylated peptides were more abundant in crotonate cultures, while the majority of butyrylated peptides were increased in butyrate cultures **(Figure 4A)**. Some β-oxidation proteins had significant abundance changes both at the protein level and at modified residues. One such case is acyl-CoA dehydrogenase (Swol_2126), which was observed in the shotgun proteomic measurements to increase in crotonate cultures; targeted PRM analyses revealed that K214 acetylation also increased significantly in crotonate grown cells **(Figure 4B)**. Some of the constitutively expressed proteins noted earlier were also found to significantly vary in acylation levels. *3-*hydroxybutyryl CoA dehydrogenase (Swol_2030) had displayed increased *3-*hydroxylbutyrylation on K58 in CM cultures. K8 of peptide STPEAMAKLAPAFK, common to K232 of acetyl-CoA transferases Swol_2051 and Swol_1934 (93% identical), showed increased acetylation and butyrylation when grown in BC **(Figure 4B)**. The increased acetylation of K232 from butyrate grown cells deviates from the trend of acetylation levels as acetylation generally was increased in crotonate-grown cells, highlighting the complexity of these modifications. That some acylation levels change significantly with cultivation condition, while others are stable, could suggest that regulatory mechanisms for lysine acylation are operating in *S. wolfei*.

**Figure 4.**
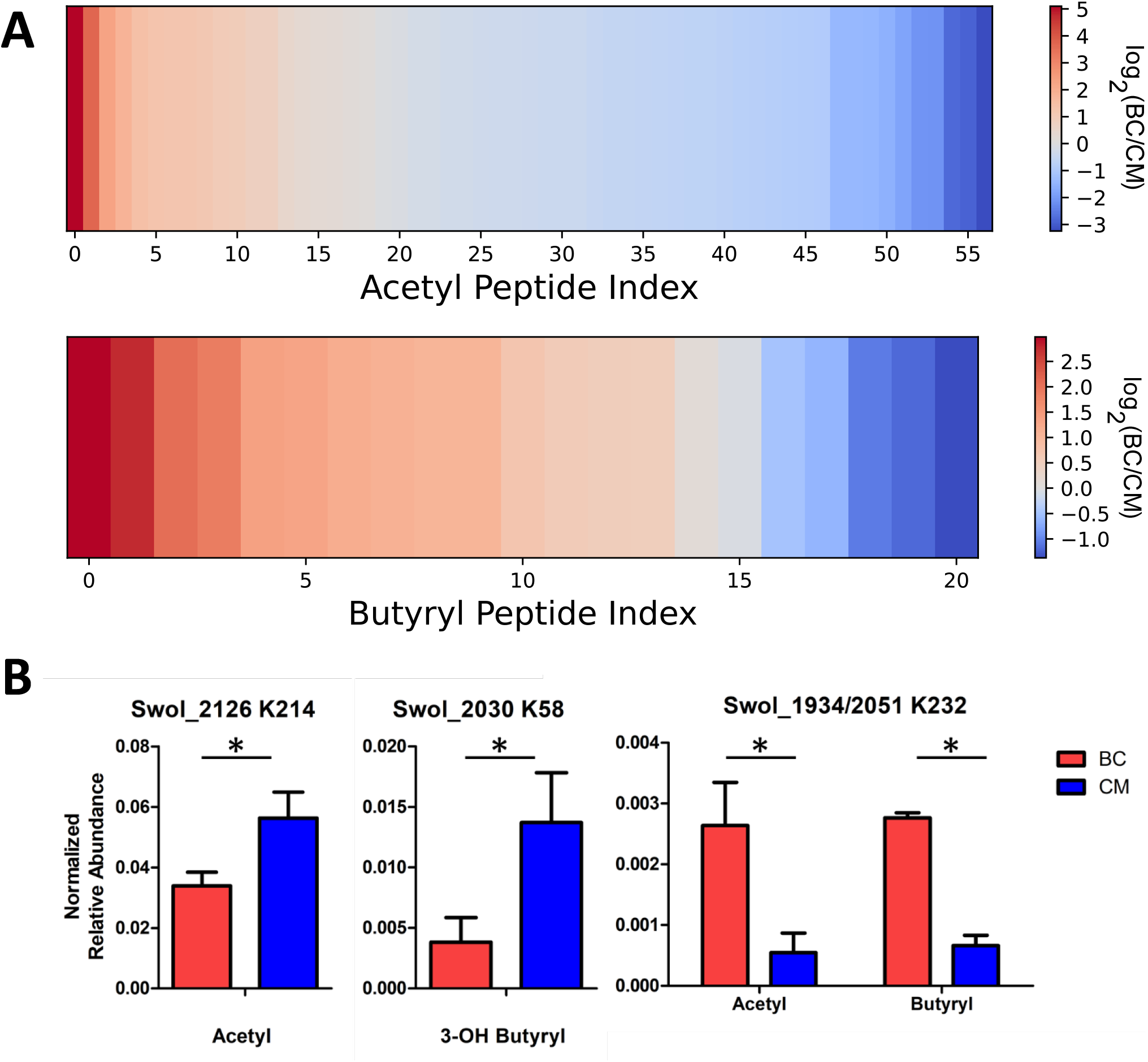
Acyl-peptide abundances vary between carbon substrates. **(A)** Heat map illustrating the fold change between growth conditions for acyl-lysine peptides involved in the butyrate degradation pathway of *S. wolfei* Göttingen. **(B)** Acyl-peptide abundances in BC and CM cultures (*p<0.05).

### 3.5 Uncovering acyl-PTM crosstalk in *S. wolfei* Göttingen

Bypassing PTM enrichment not only provides a comprehensive view of acyl-modifications proteome-wide, but also allows for the detection of peptides bearing mixed acylations and identifying PTM cross talk, which is the combination of different PTMs that can alter protein activity (Venne, Kollipara & Zahedi, 2014; Leutert, Entwisle & Villén, 2021). Many examples of intra-protein PTM crosstalk were observed, including instances of different acyl-PTMs modifying the same residue and peptides harboring multiple acyl-PTMs. In the β-oxidation pathway, 30 sites on 11 proteins were found where acyl-groups competed to modify the same residue **(Table 2)**. Enzymes with similar abundances between BC and CM cultures (Swol_0436, Swol_2052, Swol_2030, and Swol_2051) were all found with lysine residues modified by a variety of acyl-groups. The relative proportion of each type of acyl-PTM shifted between BC and CM grown cells, as illustrated by the acyl-lysine site occupancies of K78, K191, K211, K232, and K238 of acetyl-CoA transferase Swol_2051 **(Figure 5A)**. These data highlight that the type and abundance of acyl-PTMs are dynamic and fluctuate with changing metabolism.

**Figure 5.**
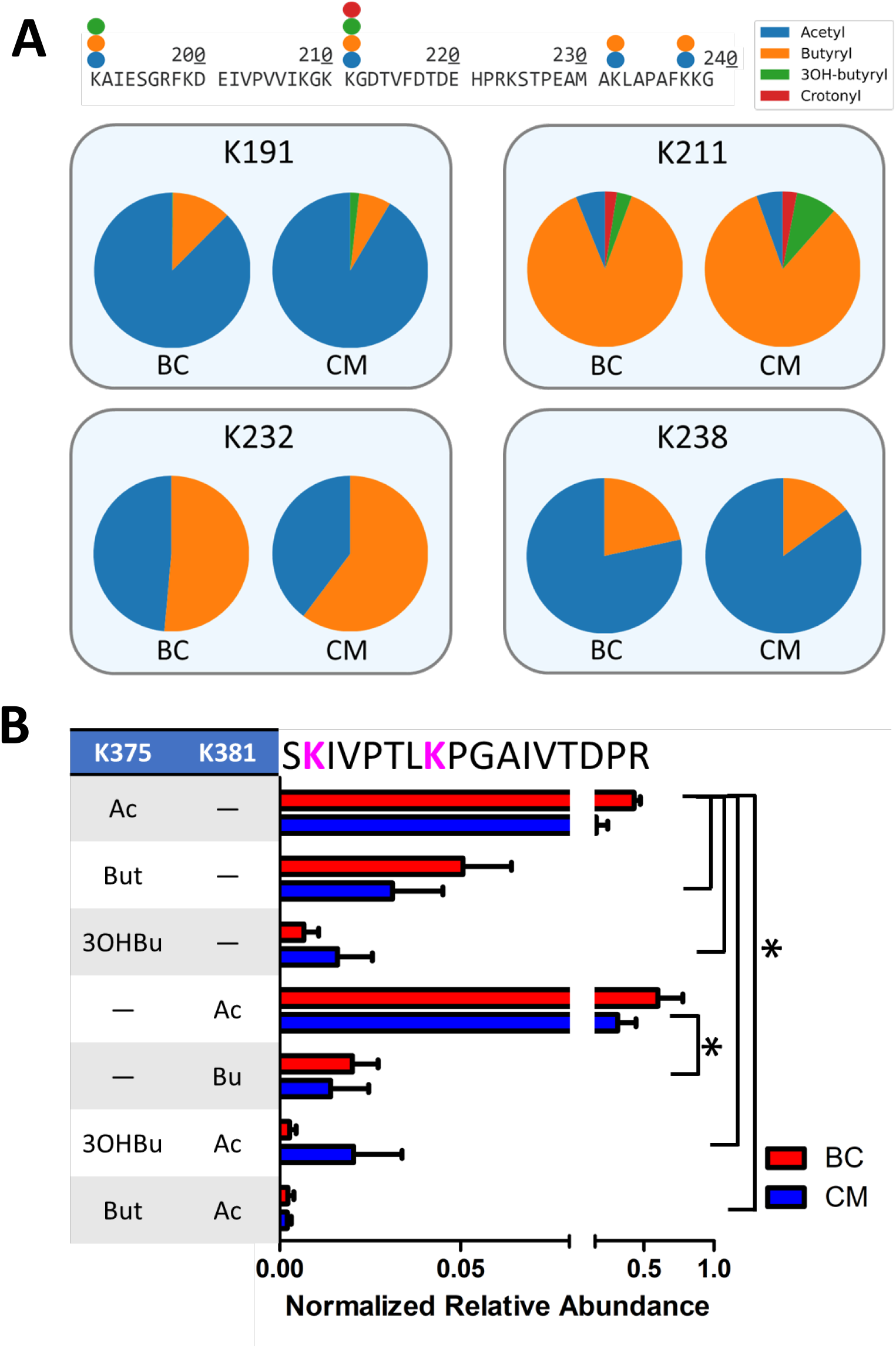
Crosstalk between acylation sites in β-oxidation enzymes. **(A)** Partial sequence (residues 190-240) of *S. wolfei* Göttingen acetyl-CoA transferase (Swol_2051) indicating sites and types of acyl-modifications. Pie charts depict relative site occupancy. **(B)** Relative abundance of different acyl combinations on peptide SKIVPTLKPGAIVTDPR (Swol_0436, residue 374-390) (*p<0.05).

Peptides containing multiple acyl modifications were also identified, potentially supporting crosstalk between proximal sites. Peptide SKIVPTLKPGAIVTDPR from acyl-CoA transferase (Swol_0436, residue 374-390) was found with 7 different acyl-PTM combinations at K375 and K381, with some versions either singly- or doubly-modified by acetyl, butyryl, or *3-*hydroxybutyryl-lysine(s). The singly-acetylated peptides, either at K375 or K381, were the most abundant out of the PTM combinations **(Figure 5B)**. Not every acylation combination was identified on K375 and K381, suggesting that there may be some regulation of these PTMs. Proteins involved in *S. wolfei* β-oxidation are generally not modified by single, isolated groups, but rather by diverse combinations that could tune enzymatic function.

### 3.6 Novel acyl-lysine modifications in *S. wolfei* subsp. *methylbutyratica* are metabolite-derived

To further explore syntrophic bacteria and acylation from RACS, we next investigated the acylome of *S. wolfei* subspecies *methylbutyratica*, a closely related syntroph that metabolizes longer carbon substrates. In addition to growth on butyrate and crotonate (C4), cells were also grown in *2-*methylbutyrate (C5), valerate (C5), and hexanoate (C6). Acyl-PTMs derived from RACS intermediates formed during substrate degradation were observed **(Supplemental Table 2)**.Collectively, 353 acylated sites on 166 proteins, were identified. The types of acyl-modifications are similar to ones identified in *S. wolfei* Göttingen and in syntroph, *S. aciditrophicus* (Muroski et al., 2022), illustrating the similarity and large number of acylations present in syntrophic bacteria **(Supplemental Table 4)**. The number and type of acyl PTMs observed across culture conditions is illustrated in **Figure 6A**.

**Figure 6.**
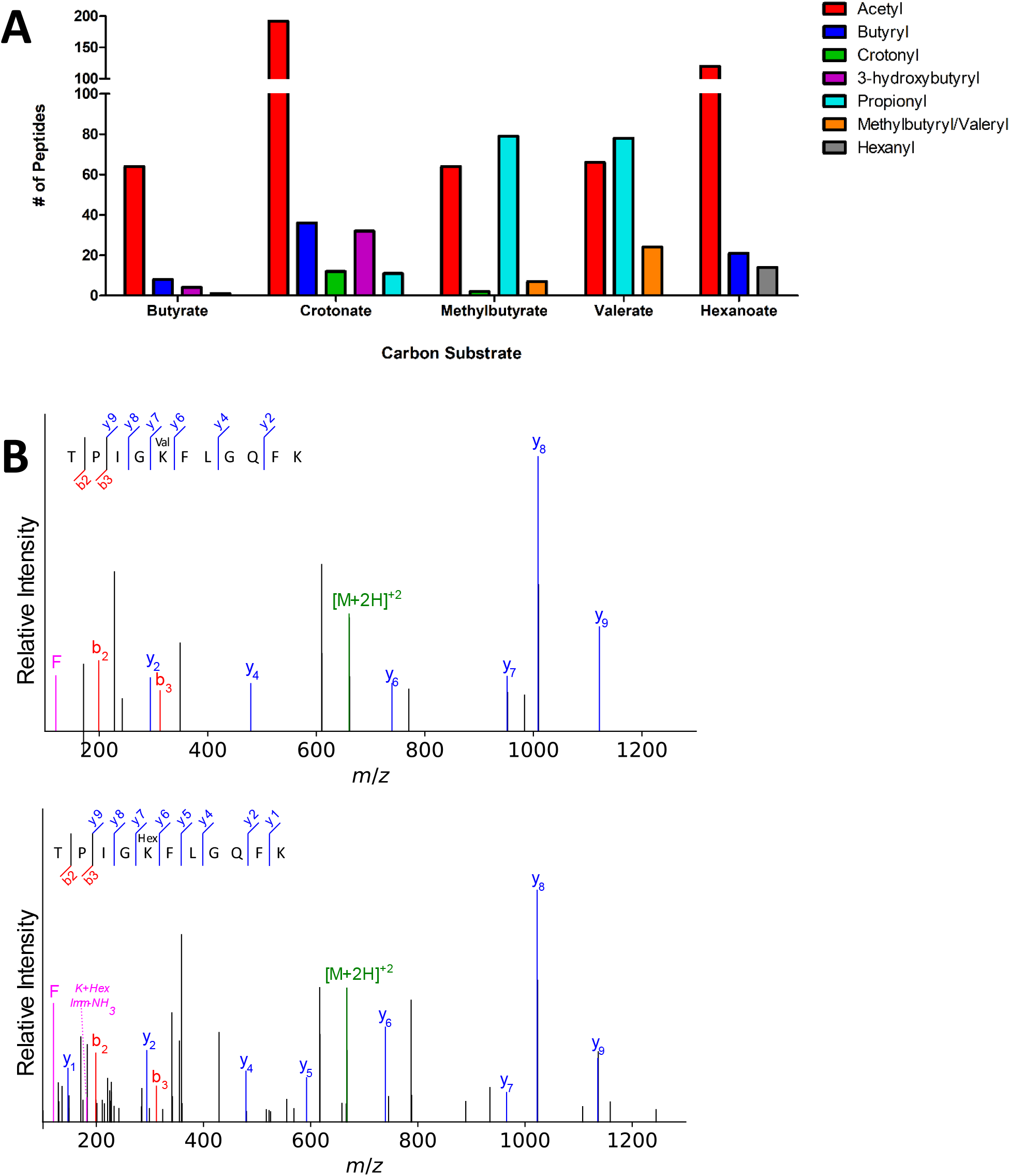
Novel acyl-lysine modifications in *S. wolfei* sub sp. *methylbutyratica*. **(A)** Acylated peptides across carbon substrates. **(B)** Tandem mass spectra of acetyl-CoA transferase (Ga0126451_1012) peptides show the 14 Da shift between valeryl- and hexanyl-lysine. Product ions (b-, y-, and immonium-related) are colored red, blue, and magenta, respectively; precursor ion is green.

In addition to the 4 types of acylation observed in *S. wolfei* Göttingen, longer acyl-modifications (methylbutyryl-/valeryl-, and hexanyl-lysine) were also observed, but only in cells grown on the corresponding carbon substrate (*i*.*e*., valeryl-lysine PTMs were observed in valerate grown cultures). Because methylbutyryl- and valeryl-are both five carbon chains (branched or linear), the two PTMs are indistinguishable by mass. They are grouped together in **Figure 6A**. The valeryl- and hexanyl-lysine modifications are the first identified in an organism. We verified the novel acyl-PTMs by comparing the tandem mass spectra and retention times to those from synthetic versions and sought acyl-lysine associated immonium ions in the tandem spectra **(Figure 6B, Supplemental Figure 3)** (Muroski et al., 2021). Propionyl-lysine modifications were primarily observed in cells grown on odd chain carbon substrates (*i*.*e*., *2-*methylbutyrate and valerate) consistent with β-oxidative degradation that would produce propionyl-CoA in addition to acetyl-CoA from odd-carbon number acyl intermediates **(Table 3, Supplemental Figure 4)**. Similarly, butyryl-lysine was not observed from cells cultivated on odd numbered carbon chain substrates and was only identified from even-carbon number substrates (butyrate, crotonate, and hexanoate) whose degradation produce butyryl-CoA or for butyrate as the substrate, make butyryl-CoA as the first step. The stark contrast in types of acyl-PTMs observed from different carbon sources suggests that the modifications directly correlate to the RACS produced by β-oxidation of the respective substrate.

**Table 3.**
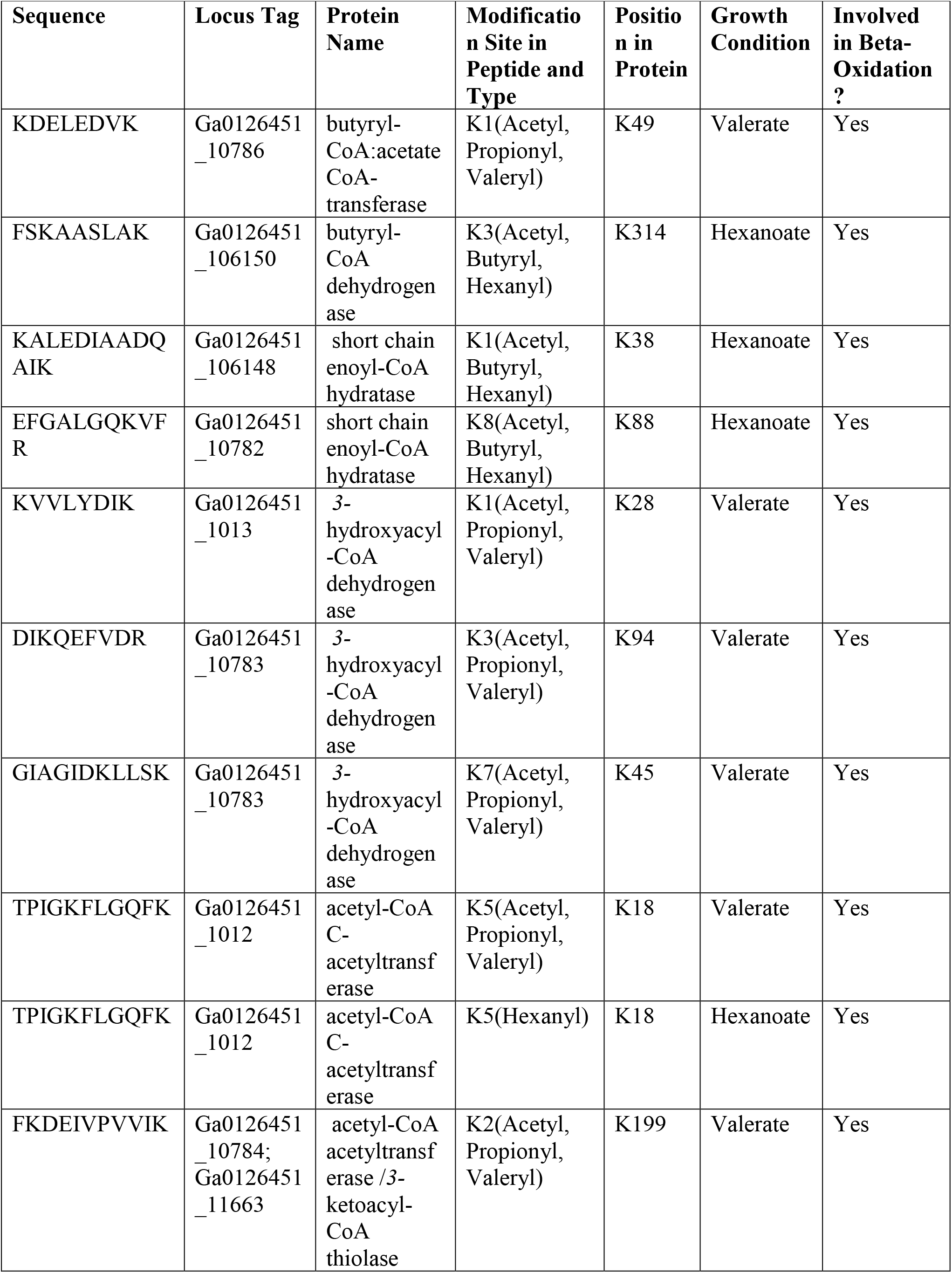
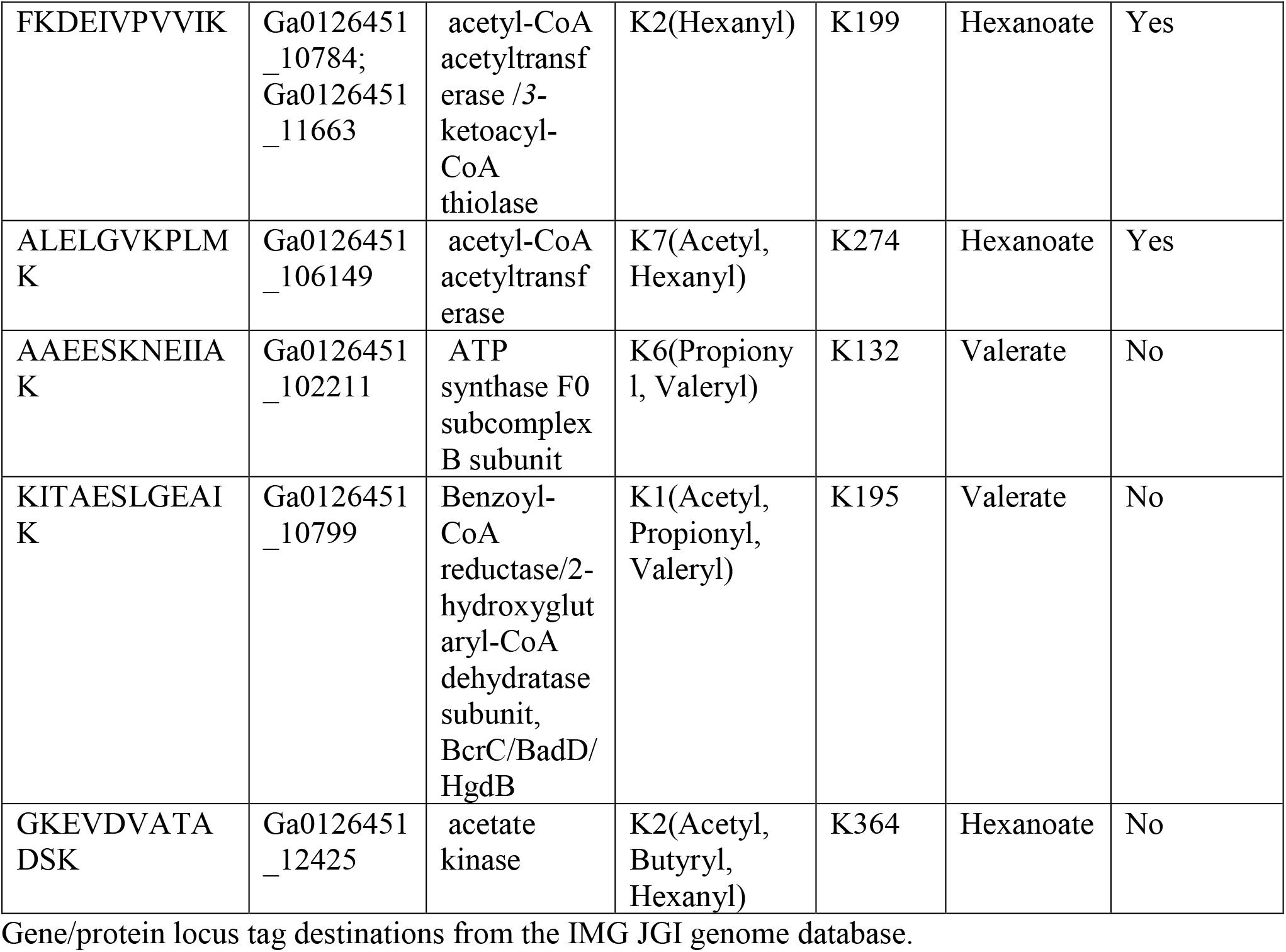
Modified peptides identified with multiple acylations in *S. wolfei* sub sp. *methylbutyratica*.

| Sequence | Locus Tag | Protein Name | Modification Site in Peptide and Type | Position in Protein | Growth Condition | Involved in Beta-Oxidation ? |
| --- | --- | --- | --- | --- | --- | --- |
| KDELEDVK | Ga0126451_10786 | butyryl-CoA:acetate CoA-transferase | K1(Acetyl, Propionyl, Valeryl) | K49 | Valerate | Yes |
| FSKAASLAK | Ga0126451_106150 | butyryl-CoA dehydrogenase | K3(Acetyl, Butyryl, Hexanyl) | K314 | Hexanoate | Yes |
| KALEDIAADQ AIK | Ga0126451_106148 | short chain enoyl-CoA hydratase | K1(Acetyl, Butyryl, Hexanyl) | K38 | Hexanoate | Yes |
| EFGALGQKVF R | Ga0126451_10782 | short chain enoyl-CoA hydratase | K8(Acetyl, Butyryl, Hexanyl) | K88 | Hexanoate | Yes |
| KVVLYDIK | Ga0126451_1013 | 3-hydroxyacyl-CoA dehydrogenase | K1(Acetyl, Propionyl, Valeryl) | K28 | Valerate | Yes |
| DIKQEFVDR | Ga0126451_10783 | 3-hydroxyacyl-CoA dehydrogenase | K3(Acetyl, Propionyl, Valeryl) | K94 | Valerate | Yes |
| GIAGIDKLLSK | Ga0126451_10783 | 3-hydroxyacyl-CoA dehydrogenase | K7(Acetyl, Propionyl, Valeryl) | K45 | Valerate | Yes |
| TPIGKFLGQFK | Ga0126451_1012 | acetyl-CoA C-acetyltransferase | K5(Acetyl, Propionyl, Valeryl) | K18 | Valerate | Yes |
| TPIGKFLGQFK | Ga0126451_1012 | acetyl-CoA C-acetyltransferase | K5(Hexanyl) | K18 | Hexanoate | Yes |
| FKDEIVPVVIK | Ga0126451_10784;<br>Ga0126451_11663 | acetyl-CoA acetyltransferase /3-ketoacyl-CoA thiolase | K2(Acetyl, Propionyl, Valeryl) | K199 | Valerate | Yes |
| FKDEIVPVVIK | Ga0126451_10784;<br>Ga0126451_11663 | acetyl-CoA acetyltransferase /3-ketoacyl-CoA thiolase | K2(Hexanyl) | K199 | Hexanoate | Yes |
| ALELGVKPLMK | Ga0126451_106149 | acetyl-CoA acetyltransferase | K7(Acetyl, Hexanyl) | K274 | Hexanoate | Yes |
| AAEESKNEIIAK | Ga0126451_102211 | ATP synthase F0 subcomplex B subunit | K6(Propionyl, Valeryl) | K132 | Valerate | No |
| KITAESLGEAIK | Ga0126451_10799 | Benzoyl-CoA reductase/2-hydroxyglutaryl-CoA dehydratase subunit, BcrC/BadD/HgdB | K1(Acetyl, Propionyl, Valeryl) | K195 | Valerate | No |
| GKEVDVATADSK | Ga0126451_12425 | acetate kinase | K2(Acetyl, Butyryl, Hexanyl) | K364 | Hexanoate | No |

Gene/protein locus tag destinations from the IMG JGI genome database.

We next investigated whether evidence of PTM crosstalk, or acyl-groups competing to modify the same residue, in metabolic enzymes was similar to that seen in the Göttingen subspecies. Several types of acyl groups modifying one residue were identified for 22 sites in 13 proteins, 10 of which are involved in β-oxidation **(Table 3)**. The three other proteins were included because they had lysine residues modified with the novel valeryl- and hexanyl-lysine modifications, as well as other acyl-PTMs. The modifications observed shifted with changes in carbon source. For example, peptide F**K**DEIVPVVIK which is common to two proteins (Ga0126451_10784; Ga0126451_11663) both of which are an acetyl-CoA acetyltransferase/ *3-*ketoacyl-CoA thiolase had different acyl groups depending on the substrate. It was modified with acetyl- and hexanyl-lysines in hexanoate-grown cells, but contained acetyl-, propionyl-, and valeryl-lysine modifications from valerate-grown cells.

This acetyl-CoA acetyltransferase is an ortholog of *S. wolfei* Göttingen Swol_2051 (95.7% sequence identity), which was also heavily acylated **(Supplemental Figure 5)**. That acylated F**K**DEIVPVVIK (K199) is common to both *S. wolfei* subsp. *methylbutyratica* and *S. wolfei* Göttingen, demonstrates that certain acylation sites are conserved **(Tables 2 and 3)**. We also observed that as the length of acyl-modification increases, so did peptide retention time, following previous reports of acylation effects on chromatography **(Supplemental Figure 6)** (Moruz et al., 2012; Mizero et al., 2021). This chromatographic behavior should be carefully considered when performing proteomic studies investigating longer chain length modifications. For valeryl- and hexanyl-lysine modifications, there were several acyl-peptides identified at retention times that corresponded to high percentages of organic phase (∼80% acetonitrile), which are commonly not analyzed in MS proteomics. Changes in modifications with carbon substrates suggest that acyl-PTMs are driven by RACS, such that the presence or absence of a modification can act as a marker for the types of RACS produced. These metabolite-driven modifications demonstrate their intimate link to the cell’s metabolic activity.

### 3.7 Crystal structure of *S. wolfei* Göttingen acetyl-CoA acetyltransferase reveals potential impacts of acylation

*S. wolfei* Göttingen uses acyl-CoA transferases to activate fatty acid substrates by synthesizing the respective CoA derivative. An acyl-CoA transferase paralog of particular interest is Act2 (Swol_0675). Acts had higher abundance levels than other Act paralogs, which did not change significantly between growth conditions. Act2 had numerous acylations and certain sites were modified with several types of acyl groups **(Table 2)**. To understand how these PTMs might affect the enzyme’s function, the structure of Act2, encoded by the gene Swol_0675, was determined by X-ray crystallography **(Figure 7A, Table 4)**. The asymmetric unit of the tetragonal crystal lattice contains two molecules of Act2, which form a homodimeric structure. About 15% of the solvent accessible surface area of the monomer, or ∼15000Å^2^, is buried in the intermolecular interface. Examination of crystal packing indicates that the Act2 homodimer likely forms a homotetramer mediated by a short β-strand-containing loop spanning residues Tyr127 to Asp140. A homotetrameric structure is consistent with the Act2 molecular mass estimated from size exclusion chromatography, with the computational prediction by the PISA webserver (Krissinel & Henrick, 2007), and with existing acetyl-CoA transferase structures that also have a similar homotetrameric structures. With this structure, we were able to visualize where the modified residues were located.

**Figure 7.**
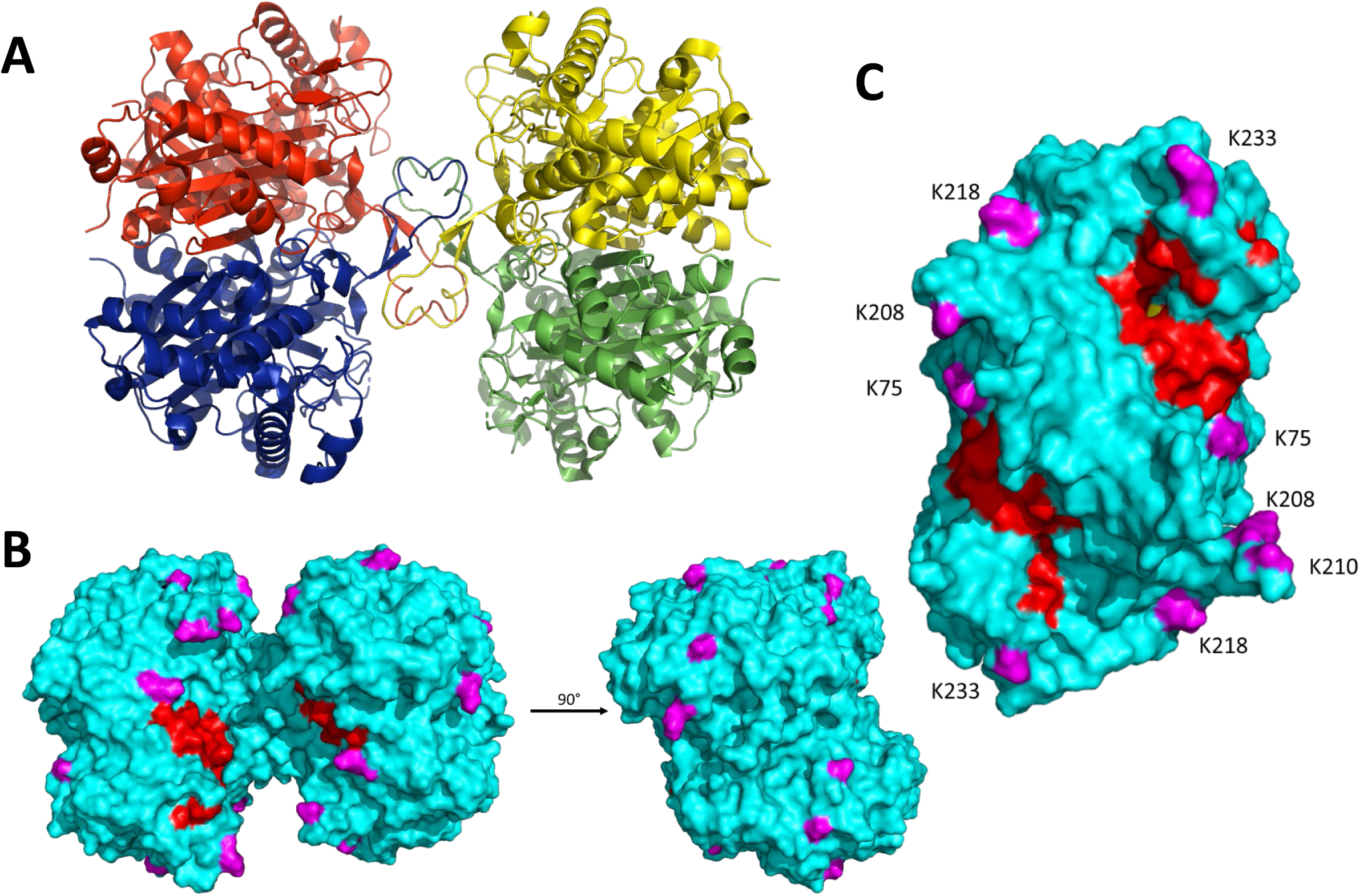
Crystal structure of *S. wolfei* Göttingen acetyl-CoA transferase, Act2 (Swol_0675; PDB entry: 7N7Z). **(A)** Act2 tetramer with monomers colored individually. **(B)** Front (same orientation as A) and side views of the molecular surface of the Act2 tetramer. Acylated residues identified from proteomic datasets are highlighted in magenta; covering and pantetheine loop are colored in red. **(C)** Side view of the molecular surface of the Act2 homodimer, with similar coloring scheme as B. Figures were generated with PyMOL (Anon, n.d.).

**Table 4.** Data collection and structural refinement statistics of *S. wolfei* Göttingen Act2.

| ***S. wolfei* Göttingen Act2 (Swol_0675)**<br>(PDB entry: 7N7Z) |  |
| --- | --- |
| **Wavelength** | 0.9792 |
| **Resolution range** | 41.64 - 2.022 (2.094 - 2.022) |
| **Space group** | P 41 21 2 |
| **Unit cell** | 81.07 81.07 242.31 90 90 90 |
| **Total reflections** | 317087 (26164) |
| **Unique reflections** | 53598 (5118) |
| **Multiplicity** | 5.9 (5.1) |
| **Completeness (%)** | 99.40 (96.93) |
| **Mean I/sigma(I)** | 10.47 (1.63) |
| **Wilson B-factor** | 33 |
| **R-merge** | 0.1041 (0.8022) |
| **R-meas** | 0.1141 (0.8933) |
| **R-pim** | 0.04584 (0.3867) |
| **CC1/2** | 0.997 (0.511) |
| **CC\*** | 0.999 (0.822) |
| **Reflections used in refinement** | 53598 (5118) |
| **Reflections used for R-free** | 5360 (512) |
| **R-work** | 0.2327 (0.3383) |
| **R-free** | 0.2743 (0.4035) |
| **CC(work)** | 0.941 (0.670) |
| **CC(free)** | 0.950 (0.445) |
| **Number of non-hydrogen atoms** | 5987 |
| **macromolecules** | 5768 |
| **ligands** | 0 |
| **solvent** | 219 |
| **Protein residues** | 795 |
| **RMS(bonds)** | 0.002 |
| **RMS(angles)** | 0.48 |
| **Ramachandran favored (%)** | 95.82 |
| **Ramachandran allowed (%)** | 3.93 |
| **Ramachandran outliers (%)** | 0.25 |
| **Rotamer outliers (%)** | 0 |
| **Clashscore** | 4.31 |
| **Average B-factor** | 42.66 |
| **macromolecules** | 42.71 |
| **solvent** | 41.43 |
| **Number of TLS groups** | 1 |

Acetyl-CoA transferases from a large variety of organisms have been structurally characterized and a search for enzymes structurally related to Act2 using the PDBeFold server (Krissinel & Henrick, 2004) allowed us to identify conserved catalytic regions in *S. wolfei* Act2 shared amongst Gram-negative and Gram-positive bacteria. The three most closely related structures are the Act structures from *Escherichia coli* (PDBid 5F38, Z-score 20.5, rmsd of 1.04 Å over 386 amino acids with sequence identity of 46%), *Clostridium difficile* (PDBid 4DD5, Z-score 19.8, rmsd of 1.05 Å over 384 amino acids with sequence identity of 48%), and *Clostridium acetobutylicum* (PDBid 4WYR, Z-score 19.7, rmsd of 1.05 Å over 386 amino acids with sequence identity of 46%) **(Supplemental Figure 7)**. Several regions are conserved among the four structures including the CoA binding region near the active site and the substrate tunnel. The position of the four key catalytic residues (C91, N319, H357, C387) is virtually identical in all four of the structures as is the shielding of the active site by the covering (residues 146-161) and pantetheine loops (residues 246-250), which were identified in the human mitochondrial acetoacetyl-CoA thiolase (T2) (Haapalainen et al., 2007) **(Supplemental Figure 8)**.

All the acyl-lysine sites identified in our proteomic datasets were mapped onto the *S. wolfei* Act2 structure and all are on the protein’s surface, with three sites located closer to the covering and pantetheine loop, K75, K208, and K233 **(Figure 7B and C)**. The position of acetylated residue K75 is conserved in the *E. coli* and *C. acetobutylicum* Act structures (substituted with glycine in *C. difficile* sequence) and the position of K233 is conserved in all four structures (although disordered in the *E. coli* structure). Residues K208 and the highly conserved K233 were found to be acetylated and/or butyrylated in *C. acetobutylicum* as found in *S. wolfei* Göttingen Act2 (Xu et al., 2018). *C. acetobutrylicum* can also generate butyryl-CoA from acetyl-CoA from homologous β-oxidation enzymes. These shared sites of modification and production of similar RACS indicates that lysine acylation may also be conserved. The proximity of these modified lysine residues near the substrate tunnel suggest that these PTMs may be connected to the local buildup of RACS in this region. While the other modification sites are located on the enzyme’s surface but not near the substrate tunnel, these lysine residues could be involved in salt-bridge interactions that mediate tetramer formation. If these interactions are present, acylation at these residues could potentially be disruptive for protein-protein interactions.

To further investigate this relationship between RACS and acyl-PTMs, we performed *in vitro* acylation assays with Act2, characterizing the sites of acylation to determine whether modified sites could be recapitulated non-enzymatically and provide insight into their potential *in vivo* regulation. Previous studies have demonstrated that *in vitro* incubation of proteins with RACS can give rise to non-enzymatic lysine acylation (Wagner & Payne, 2013; Parks & Escalante-Semerena, 2020; Baldensperger & Glomb, 2021), but the likelihood of lysine modification on a per residue basis in Act2 is largely unexplored. Factors such as local RACS concentrations and protein-protein interactions affecting site accessibility may impact lysine modification *in vivo*. Purified recombinant *S. wolfei* Göttingen Act2 was incubated either with acetyl-CoA or butyryl-CoA and lysine modifications were detected via LC-MS/MS. Out of the 25 lysine residues in Act2, 23 sites are located on the surface and are solvent-accessible. We detected 12 sites that were acylated *in vitro* and 5 of these sites matched sites that were found to be acylated in the proteomic datasets **(Table 5)**. The lysine modifications identified *in vitro* are similar to those found in *S. wolfei in vivo*, indicating that a buildup of RACS can, indeed, yield non-enzymatic lysine acylations. Out of these 5 sites, K75, K208, and K304 were located near the covering and pantetheine loops that surrounds the substrate tunnel **(Figure 7C)**, suggesting that these acyl modifications could affect substrate accessibility. We also observed lysine modifications that were either found only *in vivo* or *in vitro* which highlights the possibility of regulation of these modified sites *in vivo* **(Table 5)**. We also observed that a majority of these residues were readily modified by acetyl-CoA, but to see any butyryl modifications it required a 50-fold increase in butyryl-CoA concentration (1 μM acetyl-CoA versus 50 μM butyryl-CoA). The large differences in acyl-CoA concentrations required for *in vitro* modification indicates that these RACS have different propensities for lysine modification (Simic et al., 2015).

**Table 5.** Sites of lysine acylation in *S. wolfei* Göttingen acetyl-CoA transferase Act2 (Swol_0675).

| **Lysine Position** | **Where located?** | **Modified *in vivo*?** | **Modified *in vitro*?** |
| --- | --- | --- | --- |
| **75** | Surface | Acetyl, Butyryl, Crotonyl, 3-Hydroxybutyr | Acetyl |
| **109** | Interior | No | No |
| **141** | Tetramerization loop | No | No |
| **170** | Surface | No | No |
| **191** | Surface | Acetyl | No |
| **208** | Surface | Acetyl, Butyryl | Butyryl |
| **210** | Surface | No | Acetyl |
| **211** | Surface | No | Acetyl |
| **214** | Surface | Acetyl | Acetyl, Butyryl |
| **218** | Surface | Acetyl | Acetyl |
| **233** | Surface | Acetyl, Butyryl | No |
| **239** | Surface | No | No |
| **240** | Surface | No | No |
| **264** | Surface | No | Acetyl |
| **265** | Surface | No | Acetyl |
| **266** | Surface | No | Acetyl |
| **273** | Surface | No | Acetyl |
| **277** | Surface | No | Acetyl |
| **289** | Surface | No | No |
| **301** | Interior | No | No |
| **304** | Surface | Acetyl | Acetyl |
| **309** | Surface | No | No |
| **334** | Surface | No | No |
| **345** | Surface | No | No |
| **381** | Surface | Acetyl | No |

Full sequence of Act 2 is in Supplemental Figure S6.

## 4 Discussion

Through our systems-level characterization of lysine acylation, we demonstrated for *S. wolfei* Göttingen and subsp. *methylbutyratica* a relationship between RACS found in the β-oxidation pathway and acyl-lysine PTMs. The acylome profiles of cells grown under different carbon substrates differ not only in the protein sites modified and the type of acylation, but also the relative abundance of modifications. We also identified novel acyl-modifications (valeryl- and hexanyl-lysine) only when the cells were fed the respective substrate and observe that the acyl-PTMs identified distinctly differ and reflect the RACS produced from degrading odd and even-chained fatty acids. These changes in the proteins’ modification, especially for metabolic enzymes whose abundances remain stable between changing growth conditions, suggests that *S. wolfei* may be regulating its metabolism post-translationally. This means of metabolic regulation has been suggested in other syntrophs **(Supplemental Table 3)** (Muroski et al., 2022). As syntrophic bacteria live in energy limited conditions, these modifications could reduce protein turnover and reduce the cell’s reliance on energetically costly protein synthesis.

These modifications could act as a metabolic brake in the cell as these modifications have been shown to reduce enzymatic activity (Crosby et al., 2010). Our structural studies of *S. wolfei* Göttingen acetyl-CoA transferase Act2 indicates that there are modified residues located near catalytically critical regions of the protein, suggesting that fluctuations in modification levels may alter the activity of metabolic enzymes. Notably, many of the enzymes we found modified form complexes with themselves or other enzymes (Crable et al., 2016), which have been demonstrated to be altered by lysine acylations in other systems (El Kennani et al., 2018; Lin et al., 2015; Zhu et al., 2019). While many of these acylations are structurally similar, differing in some cases by only a methyl or hydroxyl group, these slight differences are sufficient to cause significant alterations for deacylase specificity and activity (McClure et al., 2017). These acyl-PTMs could act as an efficient mechanism for fine-tuning its catabolic pathways.

Implementing acyl-PTMs as a negative feedback inhibitor could reduce carbon and reductive stress. This means of metabolic regulation could be useful for pathways, like β-oxidation, that incorporate reversible steps. This flexibility in pathway directionality is a phenomenon observed in other syntrophic bacteria (James et al., 2019). When cellular reducing power is excessive, the cell might acylate NAD-dependent metabolic enzymes like *3-*hydroxybutyryl CoA dehydrogenase Swol_2030, which was found to be acylated, to slow the production of NADH. If NADH levels become high, this would lead to increases in H2 partial pressures, which may make further metabolism energetically difficult. If these NAD-dependent metabolic enzymes were acylated, however, both the upstream and downstream reactions would slow, until NAD^+^ could be regenerated by other means. In syntrophs, NADH reoxidation by ferredoxin-independent hydrogenases or formate dehydrogenases can only proceed if H2 partial and formate levels are low and a methanogenic partner is needed to maintain these levels (Schink, 1997; Losey et al., 2017; Agne et al., 2022). Acyl-CoA dehydrogenases Swol_2126 and _2030 had increased acylation in the crotonate monocultures, suggesting that these higher acylation levels may indeed be a response to the lack of a methanogenic partner. Furthermore, one can also envision that changes in acylation profiles in *S. wolfei* cocultures may be in response to changes in the methanogen’s metabolism. If hydrogen consumption by the methanogenic partner is altered, NADH reoxidation in syntrophs will be affected, changing the flux of its NAD-dependent metabolic pathways and resulting in a buildup of RACS.

As these metabolite driven acyl-modifications could impact metabolic activity, deacylases such as sirtuins, may regulate these PTMs. Sirtuins are NAD^+^ dependent enzymes that can remove a variety of acyl-lysine modifications and bacterial sirtuins with promiscuous deacylase activities have been identified in model systems like *E. coli* and in another syntroph, *S. aciditrophicus* (Bheda et al., 2016; Zhao, Chai & Marmorstein, 2004; Muroski et al., 2022). Sirtuins have been linked with stress resistance mechanisms related to oxidative stress and carbon starvation (Abouelfetouh et al., 2015; Ma & Wood, 2011). Sirtuin activity also depends on NAD^+^, linking lysine acylation with the cellular redox state. Sirtuins may be active in this system as the uneven distribution of acylation abundances in the proteome and *in vitro* acylation of Act2 data both suggest that there is regulation of these PTMs. In *S. wolfei* Göttingen, we identified a putative sirtuin (Swol_1033) by sequence homology that is predicted to be membrane-bound. Characterization of its substrate specificity and activity remains an important direction for future studies of this bacterium.

To further understand the metabolic impact of lysine acylation in syntrophs and other organisms, we should consider characterizing them in a comprehensive and unbiased manner. Mass spectrometry-based proteomics is a powerful tool for analyzing the acylome (Xu, Shi & Bao, 2022). Proteomics can identify the broad scope of PTMs occurring within a biological system, but analyzing a wide range of acylations can present analytical challenges such as potential sequence misidentifications (Lee et al., 2013; Kim, Zhong & Pandey, 2016). The ambiguities from isomeric and/or isobaric combinations must be considered while assigning modified residues in proteomic datasets, especially when characterizing PTM crosstalk on multiply modified peptides. Incorporated diagnostic marker ions of lysine acylation in proteomics analyses can help increase confidence in the PTMs identified, particularly when peptides contain multiple acyl modifications **(Supplemental Figure 2)** (Muroski et al., 2021).

In order to better understand how multiple acyl-PTMs relate to one another in *S. wolfei*, MS techniques such as middle-down and top-down proteomics may be useful for characterizing PTM crosstalk (Leutert, Entwisle & Villén, 2021). Unlike bottom-up proteomics, which was performed in this study and digests proteins into peptides, middle-down proteomics utilizes different digestion strategies to obtain partially digested proteins. This approach generates longer peptides, increasing the likelihood of observing co-occurring PTMs. Top-down proteomics does not incorporate any protein digestion and instead measures whole proteins. This technique provides proteoform level information, allowing one to identify PTM combinations present on a protein regardless of their sequence proximity. There remain significant challenges, however, in top-down proteomics as it is limited by low throughput, low proteome coverage, and the complexity of acylations adds great ambiguity to the data interpretation. To characterize PTM crosstalk, one must consider the combinatorial nature of these acyl modifications on multiple potential sites which generates large numbers of theoretical proteoforms for consideration and computational PTM assignments must be carefully inspected (Moradian et al., 2014; Zhou, Paša-Tolić & Stenoien, 2017). It is also possible that unknown modifications might be missed. Further investigation of these acyl-PTMS in combination with each other is important to fully understanding how these critical environmental microbial consortia regulate their metabolism. More broadly, additional studies of acyl-PTMs in other bacteria may further uncover the intricate link between RACS intermediates and its impact on metabolic regulation.

## 5 Data availability statement

Mass spectrometry data have been deposited to the ProteomeXchange Consortium (proteomexchange.org) via the MassIVE partner repository with the dataset identifier PXD034881. Crystallographic data, refinement statistics, and the structure has been deposited in the Protein Data Bank under accession code 7N7Z.

## 7 Author Contributions

JYF, JMM, MJM, RPG, RROL, JAL designed research; JYF, JMM, MAA, JAS, NQW, performed research; JYF, JMM, MAA, JAS, NQW, MJM, RROL, RPG analyzed data; and JYF, JMM, MAA, MJM, RPG, RROL, JAL wrote and edited the paper.

## 9 Abbreviations

ABC: ammonium bicarbonate
AGC: automatic gain control
BC: butyrate coculture
CoA: Coenzyme A
CM: crotonate monoculture
DDA: data-dependent acquisition
DTT: dithiothreitol
eFASP: enhanced filter-assisted sample preparation
HCD: high-energy collisional dissociation
HILIC: hydrophilic interaction chromatography
IAM: iodoacetamide
KEGG: Kyoto Encyclopedia of Genes and Genomes
LC-MS/MS: liquid chromatography tandem mass spectrometry
MS: mass spectrometry
NAD+/NADH: nicotinamide adenine dinucleotide (oxidized/reduced)
NE-CAT: Northeastern Collaborative Access Team
RACS: reactive-acyl Coenzyme A species
PRM: parallel reaction monitoring
PTMs: post-translational modifications

