## Supplementary Material for "Dynamic acylome reveals metabolite driven modifications in *Syntrophomonas wolfei*"

### 1 Supplementary Tables

**Supplemental Table 1.** Ions targeted in PRM experiments for quantitative analysis

(Please refer to attached Excel spreadsheet.)

**Supplemental Table 2.** Acyl modifications searched for in proteomic datasets

| Acyl Modification | Chemical Formula | Monoisotopic Mass Shift | Average Mass Shift | Immonium Ion | Cyclized Immonium Ion |
|---|---|---|---|---|---|
| Acetyl | $C_2H_2O$ | 42.01056 | 42.03677 | 143.1179 | 126.0913 |
| Butyryl | $C_4H_6O$ | 70.04186 | 70.09001 | 171.1492 | 154.1226 |
| Crotonyl | $C_4H_4O$ | 68.02621 | 68.07413 | 169.1336 | 152.107 |
| *3-H*ydroxybutyryl | $C_4H_6O_2$ | 86.03678 | 86.08942 | 187.1441 | 170.1176 |
| Acetoacetyl | $C_4H_4O_2$ | 84.02113 | 84.07353 | 185.1285 | 168.1019 |
| Valeryl | $C_5H_8O$ | 84.05751 | 84.11663 | 185.16475 | 168.13821 |
| Pentenoyl | $C_5H_6O$ | 82.04186 | 82.10075 | 183.14916 | 166.12256 |
| *3-H*ydroxypentanoyl | $C_5H_8O_2$ | 100.0524 | 100.116 | 201.1597 | 184.1331 |
| *3*-Oxopentanoyl | $C_5H_6O_2$ | 98.03678 | 98.10016 | 199.14408 | 182.11748 |
| Propionyl | $C_3H_4O$ | 56.0262 | 56.06339 | 157.1266 | 140.1069 |
| Hexanoyl | $C_6H_{10}O$ | 98.07316 | 98.14326 | 199.18046 | 182.15386 |
| Hexenoyl | $C_6H_8O$ | 96.05751 | 96.12737 | 197.16481 | 180.13821 |
| *3*-Hydroxy-Hexanoyl | $C_6H_{10}O_2$ | 114.0681 | 114.1427 | 215.1754 | 198.1488 |
| *3*-Oxohexanoyl | $C_6H_8O_2$ | 112.0524 | 112.1268 | 213.1597 | 196.1331 |

**Supplemental Table 3.** Proteins and peptides identified in *S. wolfei* Göttingen and *S. wolfei* sub sp. *methylbutyratica,*

(Please refer to attached Excel spreadsheet.)

**Supplemental Table 4.** Unique acyl peptides identified in proteomic studies of *S. wolfei* Göttingen*, S. wolfei* sub sp. *methylbutyratica,* and *Syntrophus aciditrophicus.*

| **# of Unique Acyl Peptides** | ***S. wolfei* Göttingen** | ***S. wolfei* sub sp. *methylbutyratica*** | ***S. aciditrophicus* SA** |
|---|---|---|---|
| Acetyl | 506 | 300 | 120 |
| Butyryl | 113 | 51 | 3 |
| *3*-Hydroxybutyryl | 55 | 34 | 6 |
| Crotonyl | 15 | 16 | 2 |
| Propionyl | *N.A.* | **123** | *N.A.* |
| Methylbutyryl/Valeryl | *N.A.* | **20** | *N.A.* |
| Glutaryl | *N.A.* | *N.A.* | **10** |
| Hexanyl | *N.A.* | **14** | *N.A.* |
| Benzoyl | *N.A.* | *N.A.* | **2** |
| *3*-Hydroxypimelyl | *N.A.* | *N.A.* | **19** |

Numbers of acyl peptides identified are summed across all carbon sources investigated in this study and in *Muroski et al.*(Muroski et al., 2021). *N.A.* denotes modifications not searched for in the proteomic datasets, because they were unrelated to RACS produced by that organism under the conditions investigated. Acyl modification types unique to the indicated strain are in bold.

### 2 Supplementary Figures

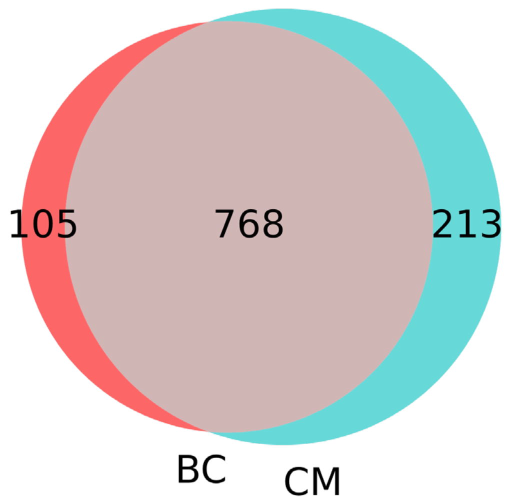


**Supplemental Figure S1. Numbers of proteins identified in butyrate cocultures (BC) and crotonate monocultures (CM) from HILIC-fractionated lysates of *S. wolfei* Göttingen.**

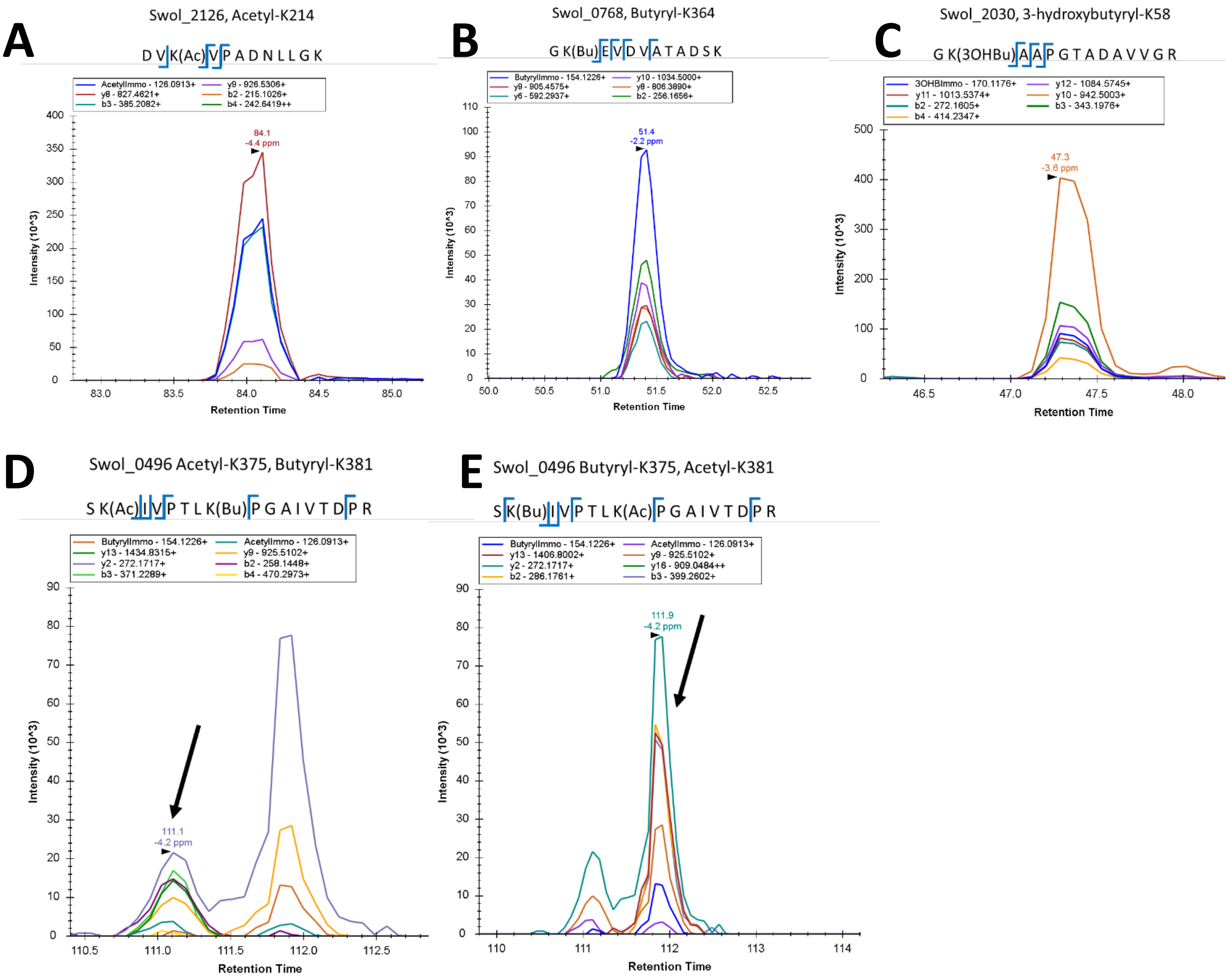


**Supplemental Figure S2. Skyline traces of MS/MS level quantitation of *S. wolfei* Göttingen acyl-peptides incorporating diagnostic immonium ions. A-C)** Extracted MS/MS ion chromatograms for DVK(Ac)VPADNLGGK (Swol_2126), GK(Bu)EVDVATADSK (Swol_0768), and GK(3OHBu)AAPGTADAVVGR (Swol_2030), respectively. Traces corresponding to each acyl-lysine immonium ion are blue. **D-E)** Elution profiles of extracted MS/MS ion chromatograms for Swol_0496 acyl-peptide isoforms SK(Ac)IVPTLK(Bu)PGAIVTDPR and SK(Bu)IVPTLK(Ac)PGAIVTDPR. Traces for the acyl-lysine immonium ions confirms that both acetyl- and butyryl-lysine modifications are present in the peptides. Retention time is in minutes.

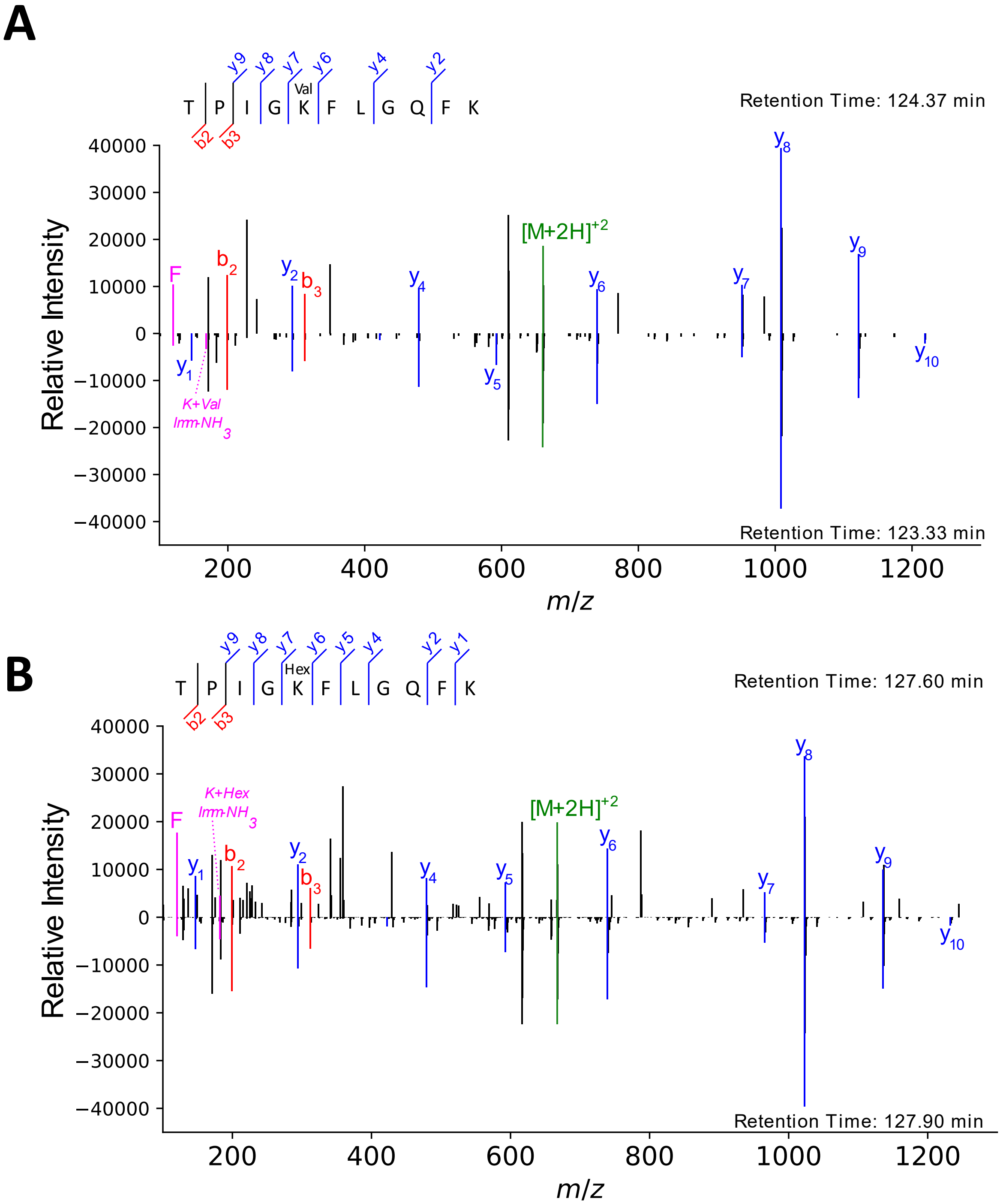


**Supplemental Figure S3. Comparing valeryl- and hexanyl-lysine modified peptides from in *S. wolfei* sub sp. *methylbutyratica* to synthetic standards. A-B)** MS/MS spectra TPIG**K**FLGQFK from acetyl-CoA transferase (Ga0126451_1012) with valeryl- and hexanyl-lysine modifications on the bolded lysine. Upper spectra are from peptides identified *in vivo* and lower spectra are from acylated reference peptides. Acyl-lysine immonium ions are highlighted in pink; b- and y-product ions are in red and blue; the precursor ion is green.

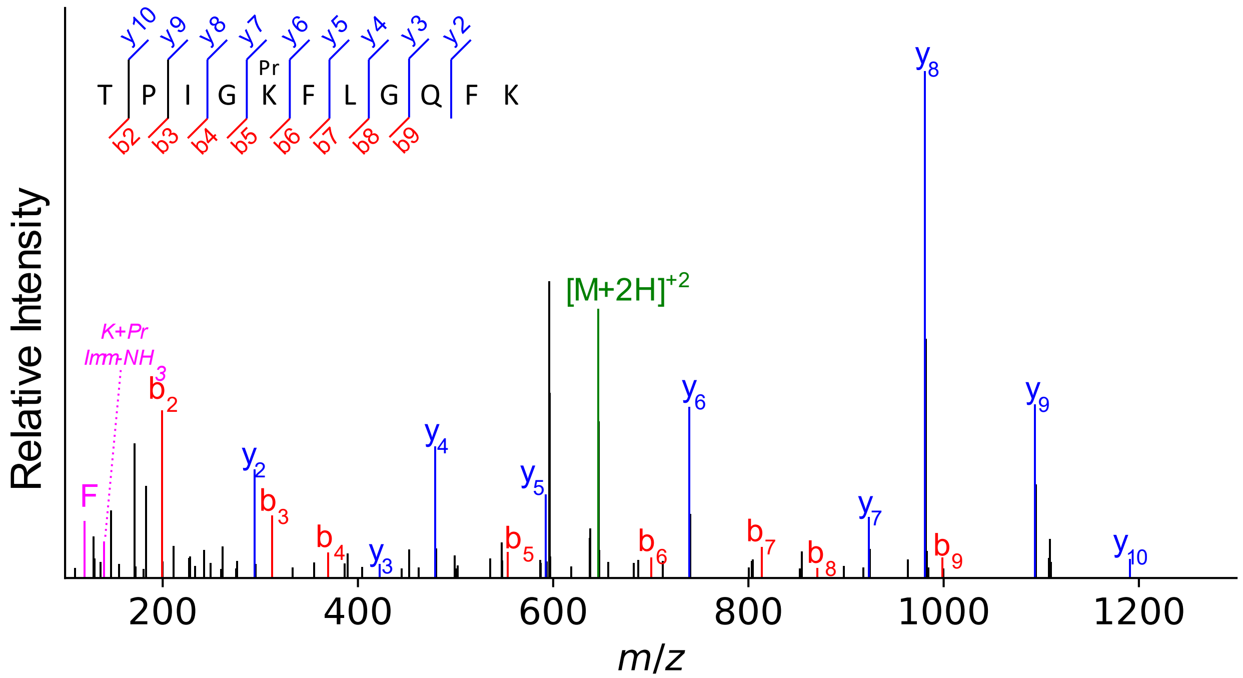


**Supplemental Figure S4. Propionyl-lysine modifications in *S. wolfei* sub sp. *methylbutyratica*.** Fragmentation spectra (MS/MS) showing mass shift corresponding to propionyl-lysine modifications on acetyl-CoA transferase (Ga0126451_1012). Product ions (b-, y-, and immonium-related) are colored in red, blue, and magenta, respectively; precursor ion is green.

```
CLUSTAL O(1.2.4) multiple sequence alignment


Ga0126451_11663      MAREVVLVGACRTPVGTFGGTIKDVGSADLGALVMGETIKRAGIKAEQIDEVIFGCVLQA      60
Swol_2051            MAREVVLVGACRTPVGTFGGTIKDVGAADLGALVMGEAITRAGIKAEQIDEVIFGCVLQA      60
                     **************************:**********:*.********************

Ga0126451_11663      GLGQNVARQCMIKAGIPKEVTAFTINKVCGSGLRAVSLAAQIIKAGDADIIMAGGTENMD      120
Swol_2051            GLGQNVARQCMIKAGIPKEITAFTINKVCGSGLRAVSLAAQVIKAGDADIILAGGTENMD      120
                     *******************:*********************:*********:********

Ga0126451_11663      KAPFLLPNARWGYRMSMPKGDLVDEMVFGGLFDIFNGYHMGITAENVNEMYGITREEQDA      180
Swol_2051            KAPFLLPNARWGYRMSMPKGDLIDEMVFGGLTDIFNGYHMGITAENVNEMYGITREEQDA      180
                     **********************:******** ****************************

Ga0126451_11663      FGFRSQELAAKAIESGRFKDEIVPVVIKGKKGDIVFDTDEHPRKSTPEAMAKLAPAFKKG      240
Swol_2051            FGFRSQDLAAKAIESGRFKDEIVPVVIKGKKGDTVFDTDEHPRKSTPEAMAKLAPAFKKG      240
                     ******:************************* ***************************

Ga0126451_11663      GSVTAGNASGINDGAAAVIVMSKEKADELGIKPMAKVVSYASGGVDPSVMGLGPIPASRK      300
Swol_2051            GSVTAGNASGINDGAAAVIVMSKEKADELGIKPMAKVVSYASGGVDPSVMGLGPIPASRK      300
                     ************************************************************

Ga0126451_11663      ALEKAGLTVDDIDLIEANEAFAAQSIAVGRDLGWSDKMEKVNVNGGAIALGHPIGASGAR      360
Swol_2051            ALEKAGLTIDDIDLIEANEAFAAQAIAVGRDLGWSDKMEKVNVNGGAIAIGHPIGASGAR      360
                     ********:***************:************************:**********

Ga0126451_11663      ILVTLLYEMQKRGSKKGLATLCIGGGQGTALIVEAI      396
Swol_2051            ILITLLYEMQKRGAKRGLATLCIGGGMGTALIVEAI      396
                     **:**********:*:********** *********
```

**Supplemental Figure S5. CLUSTAL Sequence alignment of acetyl-CoA acetyltransferases from *S. wolfei* sub sp. *methylbutyratica* and *S. wolfei* Göttingen.** Alignment of *S. wolfei* sub sp. *methylbutyratica* (Ga0126451_1153) to *S. wolfei* Göttingen Swol_2051. There is 95.7% sequence identity. The acylated peptide spanning residues 198-208, F**K**DEIVPVVIK (AcK199), identified in proteomic data of both *S. wolfei* sub species is highlighted in yellow.

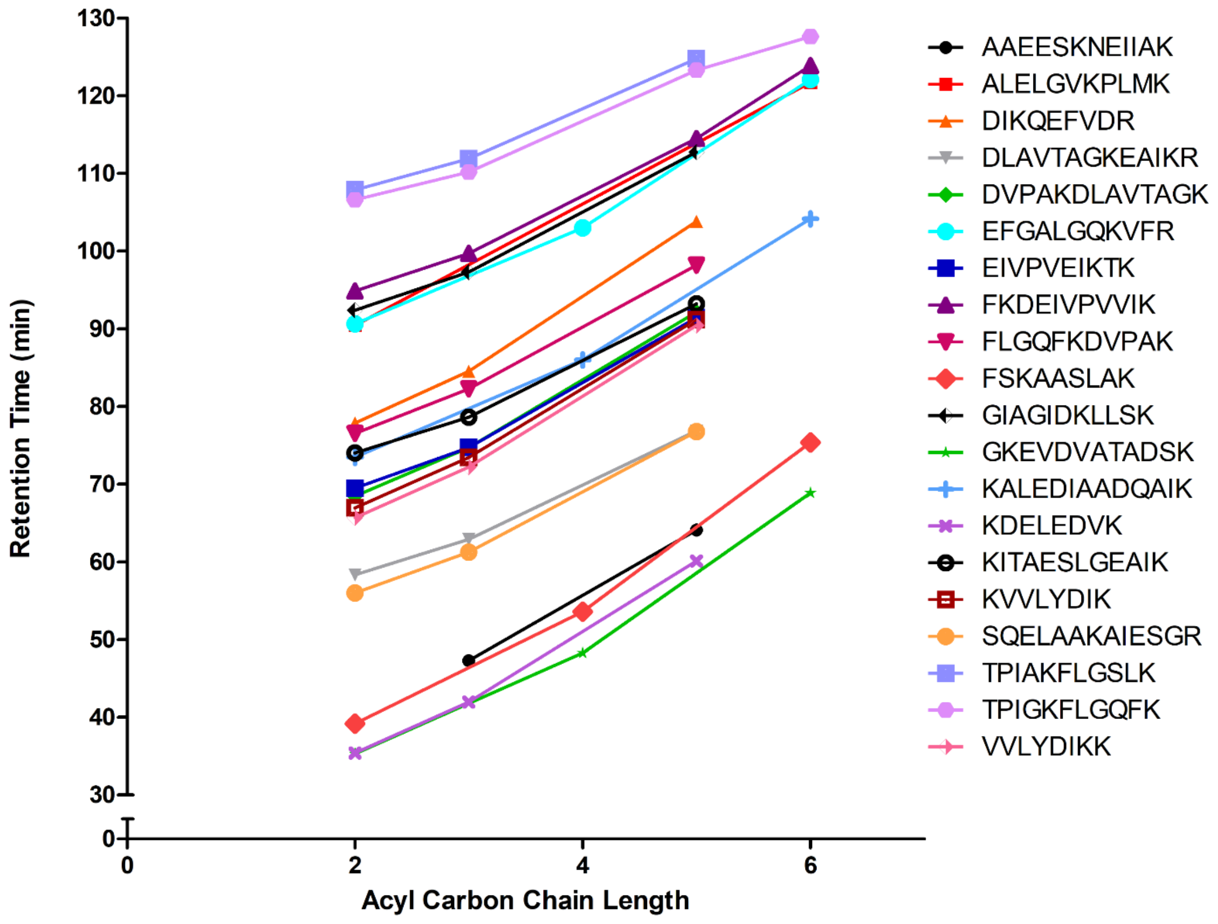


**Supplemental Figure S6. Comparison of acyl-peptide liquid chromatography retention time *versus* acyl-PTM carbon chain length.** Twenty acyl-peptides from *S. wolfei* sub sp. *methylbutyratica* were found to be modified with at most 4 types of acyl-groups. Carbon chain lengths of 2, 3, 4, 5, and 6 correspond to acetyl-, propionyl-, butyryl-, valeryl-, and hexanyl-lysine modifications.

```
CLUSTAL O(1.2.4) multiple sequence alignment


tr|Q0AZ52|Q0AZ52_SYNWW      -MAEINEVVVVGMARTPIGRYLGGLASVRANDLAIIAANAAIERAGVDPGIIDEIVGATC      59
sp|Q9BWD1|THIC_HUMAN        MNAGSDPVVIVSAARTIIGSFNGALAAVPVQDLGSTVIKEVLKRATVAPEDVSEVIFGHV      60
sp|P76461|ATOB_ECOLI        ----MKNCVIVSAVRTAIGSFNGSLASTSAIDLGATVIKAAIERAKIDSQHVDEVIMGNV      56
sp|Q18AR0|THLA_CLOD6        ----MREVVIASAARTAVGSFGGAFKSVSAVELGVTAAKEAIKRANITPDMIDESLLGGV      56
sp|P45359|THLA_CLOAB        ----MKEVVIASAVRTAIGSYGKSLKDVPAVDLGATAIKEAVKKAGIKPEDVNEVILGNV      56
                                    *:.. .** :* :  .:  . . :*.  . : .:::* :    :.* : .

tr|Q0AZ52|Q0AZ52_SYNWW      LHAGNGSLPPRIIGMKVGLPVRSGSCMVSQNCASGMRATEIACQNIMLGKTDISLVTAVE      119
sp|Q9BWD1|THIC_HUMAN        LAAGCGQNPVRQASVGAGIPYSVPAWSCQMICGSGLKAVCLAVQSIGIGDSSIVVAGGME      120
sp|P76461|ATOB_ECOLI        LQAGLGQNPARQALLKSGLAETVCGFTVNKVCGSGLKSVALAAQAIQAGQAQSIVAGGME      116
sp|Q18AR0|THLA_CLOD6        LTAGLGQNIARQIALGAGIPVEKPAMTINIVCGSGLRSVSMASQLIALGDADIMLVGGAE      116
sp|P45359|THLA_CLOAB        LQAGLGQNPARQASFKAGLPVEIPAMTINKVCGSGLRTVSLAAQIIKAGDADVIIAGGME      116
                            * ** *.   *    . *:     .   .  *.**:::. :* * *  *.:.  :. . *

tr|Q0AZ52|Q0AZ52_SYNWW      SMSNIPYLLQ-QARSGYRMGDGKVQDAMLSDGLVCQLAGGHMGMTAENIAEKYGITREEC      178
sp|Q9BWD1|THIC_HUMAN        NMSKAPHLAY--LRTGVKIGEMPLTDSILCDGLTDAFHNCHMGITAENVAKKWQVSREDQ      178
sp|P76461|ATOB_ECOLI        NMSLAPYLLDAKARSGYRLGDGQVYDVILRDGLMCATHGYHMGITAENVAKEYGITREMQ      176
sp|Q18AR0|THLA_CLOD6        NMSMSPYLVP-SARYGARMGDAAFVDSMIKDGLSDIFNNYHMGITAENIAEQWNITREEQ      175
sp|P45359|THLA_CLOAB        NMSRAPYLAN-NARWGYRMGNAKFVDEMITDGLWDAFNDYHMGITAENIAERWNISREEQ      175
                            .**  *:*     * * ::*:  . * :: ***     . ***:****:*:.: ::**

tr|Q0AZ52|Q0AZ52_SYNWW      DALALTSHQNAVKAVDEGIFDREIVPVVIKSKKGDKVISKDEHPIRGASLETMAKLPPAF      238
sp|Q9BWD1|THIC_HUMAN        DKVAVLSQNRTENAQKAGHFDKEIVPVLVSTRKGLIEVKTDEFPRHGSNIEAMSKLKPYF      238
sp|P76461|ATOB_ECOLI        DELALHSQRKAAAAIESGAFTAEIVPVNVVTRKKTFVFSQDEFPKANSTAEALGALRPAF      236
sp|Q18AR0|THLA_CLOD6        DELALASQNKAEKAQAEGKFDEEIVPVVIKGRKGDTVVDKDEYIKPGTTMEKLAKLRPAF      235
sp|P45359|THLA_CLOAB        DEFALASQKKAEEAIKSGQFKDEIVPVVIKGRKGETVVDTDEHPRFGSTIEGLAKLKPAF      235
                            * .*: *:..:  *   * *  ***** :  :*    .. **.   .:. * :. * * *

tr|Q0AZ52|Q0AZ52_SYNWW      KKGG--VVTAANASGINDCAAAAVFMSKKKCEELGLKPLMKLVGICSEGVDAKVMGLGPA      296
sp|Q9BWD1|THIC_HUMAN        LTDGTGTVTPANASGINDGAAAVVLMKKSEADKRGLTPLARIVSWSQVGVEPSIMGIGPI      298
sp|P76461|ATOB_ECOLI        DKAG--TVTAGNASGINDGAAALVIMEESAALAAGLTPLARIKSYASGGVPPALMGMGPV      294
sp|Q18AR0|THLA_CLOD6        KKDG--TVTAGNASGINDGAAMLVVMAKEKAEELGIEPLATIVSYGTAGVDPKIMGYGPV      293
sp|P45359|THLA_CLOAB        KKDG--TVTAGNASGLNDCAAVLVIMSAEKAKELGVKPLAKIVSYGSAGVDPAIMGYGPF      293
                             . *  .** .****:** **  *.*  . .   *: **  : .    **   :** **

tr|Q0AZ52|Q0AZ52_SYNWW      VAMPKVLKQAGWKYEDVDYWEVNEAFAAQVLGVVRMLKEEAGIELDFSKTNHNGSGIGLG      356
sp|Q9BWD1|THIC_HUMAN        PAIKQAVTKAGWSLEDVDIFEINEAFAAVSAAIVKEL------GLNPEKVNIEGGAIALG      352
sp|P76461|ATOB_ECOLI        PATQKALQLAGLQLADIDLIEANEAFAAQFLAVGKNL------GFDSEKVNVNGGAIALG      348
sp|Q18AR0|THLA_CLOD6        PATKKALEAANMTIEDIDLVEANEAFAAQSVAVIRDL------NIDMNKVNVNGGAIAIG      347
sp|P45359|THLA_CLOAB        YATKAAIEKAGWTVDELDLIESNEAFAAQSLAVAKDL------KFDMNKVNVNGGAIALG      347
                             *   .:  *.    ::*  * ******   .: : *       :: .*.* :*..*.:*

tr|Q0AZ52|Q0AZ52_SYNWW      HPVGATGLRIIVSMYYELERLGLTKGGASLCVGGGSAMASLWTRDI      402
sp|Q9BWD1|THIC_HUMAN        HPLGASGCRILVTLLHTLERMGRSRGVAALCIGGGMGIAMCVQRE-      397
sp|P76461|ATOB_ECOLI        HPIGASGARILVTLLHAMQARDKTLGLATLCIGGGQGIAMVIERLN      394
sp|Q18AR0|THLA_CLOD6        HPIGCSGARILTTLLYEMKRRDAKTGLATLCIGGGMGTTLIVKR--      391
sp|P45359|THLA_CLOAB        HPIGASGARILVTLVHAMQKRDAKKGLATLCIGGGQGTAILLEKC-      392
                            **:*.:* **:.:: : ::  . . * *:**:***  . :    :
```

**Supplemental Figure S7. CLUSTAL Sequence alignment of acetyl-CoA transferases from model organisms to *S. wolfei* Göttingen.** Alignment with *S. wolfei* Göttingen Swol_0675, *Homo sapiens* ACAT2 (Q9BWD1), and *Escherichia coli* atoB (P76461), *Clostridium difficile* (Q18AR0), and *C. acetobutylicum* (P45359). Catalytic residues from the four active site loops are highlighted in yellow; covering loop and pantetheine loop are highlighted in blue. Acylated residues identified in *S. wolfei* Göttingen are highlighted in magenta; conserved sites of acylation between *S. wolfei* Göttingen and *C. acetobutylicum* are marked green. Symbols are (:) identical and (*) similar amino acids.

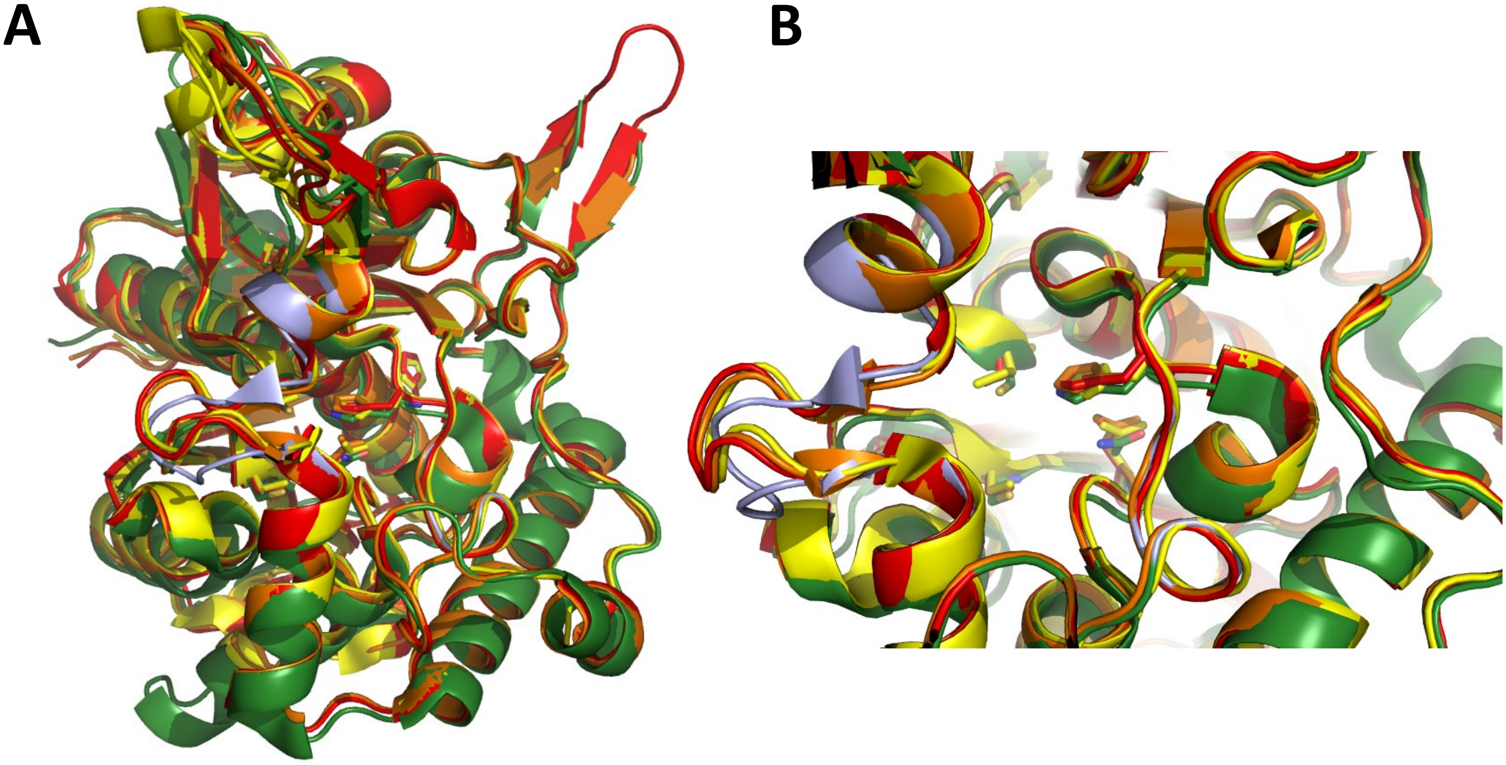


**Supplemental Figure S8. Overlay of acetyl-CoA acetyltransferase structures from *S. wolfei* Göttingen and other related homologs in Gram-positive and Gram-negative bacteria.** Structural alignment of Act structures from *S. wolfei* Göttingen (green; PDBid 7N7Z), *E. coli* (yellow; PDBid 5F38), *Clostridium difficile* (orange; PDBid 4DD5), and *C. acetobutylicum* (red; PDBid 4WYR). Catalytic loops are in light blue. **(A)** Overall structure of bacterial Acts. **(B)** Active site of Acts.
